# From naïve pluripotency to human neural organoids through a three-dimensional morphogenetic continuum

**DOI:** 10.1101/2024.11.14.623407

**Authors:** Cecilia Laterza, Elisa Cesare, Hannah T. Stuart, Martina D’Ercole, Alessia Gesualdo, Maria Grazia La Barbera, Sara Brignani, Carmela Ribecco, Roberta Polli, Roberta Frison, Silvia Angiolillo, Andrea Maset, Onelia Gagliano, Davide Cacchiarelli, James Briscoe, Elly M. Tanaka, Alessandra Murgia, Nicola Elvassore

## Abstract

Human central nervous system (CNS) development involves complex transitions from pluripotency to regionalised neural tissues. The early phases of this process are inaccessible in humans but can potentially be modelled *in vitro* using brain organoids, including to study neurodevelopmental disorders. However, current methods are based on post-implantation-like human pluripotent stem cells (hPSCs), which exhibit a hypermethylated state and show epigenetic memory retention. Here we developed a 3D model of human CNS development, starting from naïve human induced PSCs (hiPSCs), which exhibit a hypomethylated pre-implantation-like state of pluripotency and develop into 3D neuroepithelial cysts in a timely morphogenetic continuum. Upon treatment with appropriate signalling cues, naïve-derived neuroepithelial cysts can be specified toward different axial identities. Extended culture of anterior-specified organoids results in forebrain-like structures containing both dorsal and ventral neural precursors as well as mature neurons, exhibiting appropriate cellular diversity and functional properties. We applied this system to model Fragile X Syndrome (FXS), an epigenetically regulated neurodevelopmental disorder. We found that FXS patient-derived naïve hiPSCs, initially demethylated at the *Fmr1* locus, gradually underwent remethylation during organoid development. In addition, *Fmr1* silencing started much earlier than can be detected by pre-natal analysis, and is concomitant with the development of mosaicisms. Our approach provides a new platform for studying human CNS development, including early epigenetic events and regional patterning, demonstrating the potential of naïve hiPSC-derived organoids for modelling neurodevelopmental disorders with complex epigenetic regulation.

**Highlights:**

- single naïve hiPSCs differentiate into 3D neuroepithelial cysts in a timely morphogenetic continuum
- signalling cues at appropriate developmental transitions can direct naïve hiPSC-derived organoids to different regional identities of the human CNS
- naïve hiPSC-derived forebrain organoids display cellular complexity representing both dorsal and ventral identities
- forebrain organoids from Fragile X Syndrome patients recapitulate the genetic instability and epigenetic dysregulation of *Fmr1* locus.

## Introduction

Development of the human central nervous system (CNS) is a continuous process involving cell identity transitions coupled with three-dimensional (3D) tissue organisation and patterning. After blastocyst implantation, pluripotent epiblast cells undergo gastrulation generating, among others, the neuroepithelial lineage^1,2^. The neuroepithelium folds and closes forming the neural tube, which is concomitantly patterned along the rostral-caudal and dorso-ventral axes. The rostral-most regions form first from the anterior epiblast and underpin brain formation, whilst the more caudal spinal cord is laid down progressively as neuromesodermal progenitors (NMPs)^3^ in the tailbud of human embryos commit to the neural lineage during posterior axis elongation^4^. Neural organoid technology provides a tool to model, *in vitro*, the early inaccessible stages of human CNS development, both in physiological and pathological contexts^5,6^. However, current methods are based on aggregates of human primed pluripotent stem cells (hPSCs), which resemble the late post-implantation epiblast stage *in vivo* and exhibit a hypermethylated state. When generated through reprogramming, human induced pluripotent stem cells (hiPSCs) retain partial epigenetic memory of the source cells, which can affect their differentiation potential and limit their reliability in patient-specific disease modelling^7–10^. Additionally, the hypermethylated state of primed hiPSCs restricts their use in modelling epigenetic landscapes and dynamics during the development of epigenetically regulated diseases, such as Fragile X Syndrome (FXS)^11^.

In this study, we envision that 3D-cultured naïve hiPSCs, the *in vitro* counterpart of the *in vivo* pre-implantation epiblast, can be used as a starting point to enter the continuum morphogenetic process toward regionalised brain organoids. The naïve identity offers an epigenetic *tabula rasa* due to its hypomethylated genome^12,13^. Indeed, a recent study showed that a transient transition toward the naïve state during somatic cell reprogramming is sufficient to erase epigenetic memory and re-establish the full differentiation potential of primed hiPSCs^14^. However, the direct differentiation of naïve hiPSCs to embryonic lineages has been challenging and plagued by low efficiencies and protracted differentiation timelines^15–18^. Recent studies have, instead, focused on understanding how to prepare naïve hiPSCs for post-implantation decisions via capacitation in 2D cultures^15^, or on harnessing naïve hiPSC potential to generate extra-embryonic lineages to build 3D pre- and peri-implantation synthetic embryo models such as blastoids and bilaminar disk models^19–21^. We recently demonstrated that, in addition to supporting long-term feeder-free self-renewal, a 3D extracellular matrix (ECM)-rich environment allows differentiation and 3D morphogenesis of naïve hiPSCs through timely capacitation (day 5) and subsequent formation of pseudostratified primed epiblast cysts (day 8)^22^. However, the full potential of naïve hiPSCs to generate long-term 3D differentiation of embryonic lineages, including CNS, decoupled from extra-embryonic lineages still remains unexplored.

Here we demonstrate that naïve hiPSCs in 3D ECM-rich environment can promptly initiate a continuous morphogenetic process towards embryonic neuroectoderm. Single naïve hiPSCs seeded in 3D undergo differentiation and morphogenesis to form an epiblast cyst, and subsequently a neuroepithelial cyst. These neuroepithelial cysts have the potential to regionalise along the anterior-posterior axis, and to differentiate into forebrain organoids containing both dorsal and ventral neural precursors, as well as mature neurons.

The development of brain organoids from hypomethylated naïve iPSCs has allowed us to address a long-standing question in FXS pathogenesis: the timing of *Fmr1* epigenetic silencing during early embryonic development. FXS is an epigenetically regulated neurodevelopmental disorder and the leading genetic cause of mental retardation and autism spectrum disorder^23^. In FXS patients, aberrant expansions of CGG trinucleotide repeats at the 5’ UTR of the *Fmr1* locus exceeding 200 repeats are classified as full mutations, and undergo epigenetic silencing through hypermethylation^11,24^. This results in the loss of the gene product Fragile X Mental Retardation Protein (FMRP), which is important for healthy brain development^25^. In contrast, expansions between 55 and 200 CGG repeats are considered pre-mutations; these do not undergo methylation and are not silenced, but result in lower FMRP levels^26–28^. Recent studies have revealed that most FXS patients (∼40%) exhibit a mosaic pattern, with a combination of pre-mutated and fully mutated alleles^29^.

Silencing of the fully mutated *Fmr1* allele appears to occur during the first trimester of gestation, as indicated by analyses of chorionic villi biopsy samples^30–33^. However, the precise timing of *Fmr1* silencing in the CNS remains uncertain, as these analyses are based on extraembryonic tissues, which display slower methylation dynamics compared to their embryonic counterparts^34^. In animal models, *Fmr1* knockouts are commonly used, since knock-in models of *Fmr1* expansions exceeding 200 repeats do not undergo methylation in mice and can not recapitulate FMRP deficiency^35^. In contrast, human models are often derived from hiPSCs with aberrant hypermethylation^36,37^, or from human embryonic stem cells (hESCs) that exhibit variable levels of methylation^38,39^. To our knowledge, none of the existing models fully recapitulate the *Fmr1* silencing process.

Here, we demonstrate that inducing a naïve pluripotent state in FXS cells can restore *Fmr1* expression, consistent with previous findings^40,41^, while preserving the CGG trinucleotide repeat expansion within the 5’ UTR of the *Fmr1* locus. From this starting point, we show that during differentiation into naïve-derived brain organoids, the fully mutated *Fmr1* allele progressively acquires methylation. Interestingly, we observe genomic instability in the unmethylated allele *in vitro*, which may correlate with the mosaicism commonly observed in FXS patients^29^. Furthermore, we demonstrate that *Fmr1* undergoes dynamic epigenetic and genetic dysregulation, which leads to the gradual suppression of FMRP production. The generation of naïve-derived brain organoids thus opens new avenues for studying early epigenetic events in human CNS development, both under physiological conditions and in disease contexts, as well as the molecular processes driving regional patterning within a 3D morphogenic continuum.

## Results

### Single naïve iPSCs can undergo 3D morphogenesis into neuroepithelial cysts

We previously reported timely developmental morphogenesis of 3D-cultured naïve hiPSCs in ECM-rich environment, toward the primed epiblast state, through the modulation of the signalling environment^22^. Here we further explored their potential to model morphogenetic processes beyond epiblast development and gastrulation, by challenging our 3D culture system to recapitulate neural induction, starting directly from single naïve hiPSCs (**Fig. 1A**). Single naïve hiPSCs seeded in 3D Matrigel were stimulated with short-term FGF2 treatment to encourage naïve exit^22^ and, simultaneously, with dual SMAD inhibition (BMP and TGFβ/NODAL pathway inhibition), which promotes neural induction from conventional hPSCs^42^ (**Fig. 1A**). After 4 days, we observed formation of cysts with a single lumen (**Fig. 1B**) and, as differentiation proceeded, we observed thickening of the epithelium (**Fig. 1B**).

**Figure 1.**
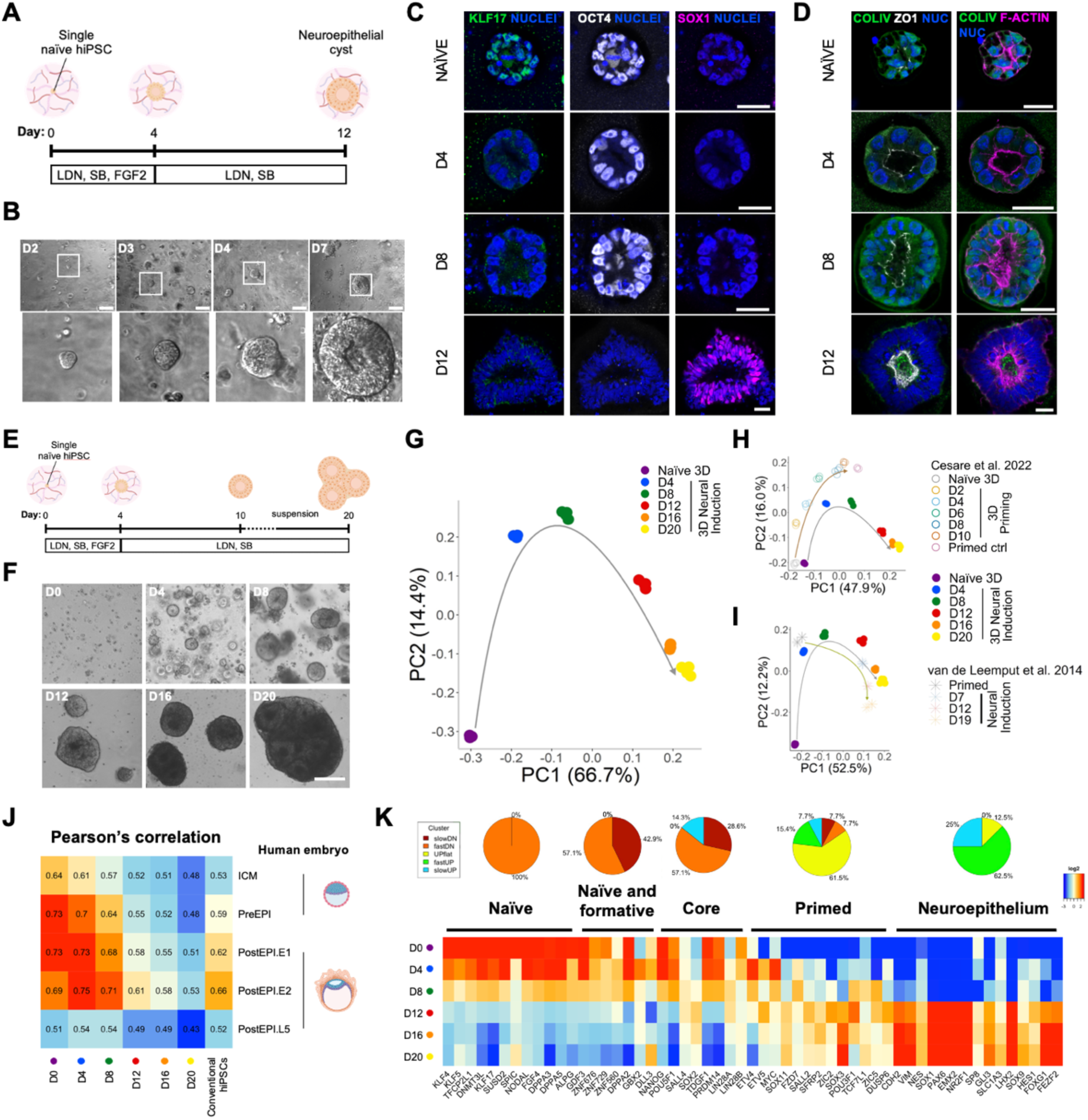
Naïve hiPSCs differentiation into neuroepithelial cysts in a 3D ECM-rich environment. **A**. Experimental design of the induction protocol. **B**. Bright field images showing the clonal growth of differentiating 3D naïve hiPSCs at D2, D3, D4 and D7. Scale bar 100 µm. **C**. Representative single confocal slices of 3D naïve, D4, D8 and D12 3D neural differentiation immunostained for KLF17 (green), OCT4 (grey) and SOX1 (magenta). Nuclei were stained with DAPI (blue). Scale bar 25 μm. **D**. Representative single confocal slices of 3D naïve, D4, D8 and D12 3D neural differentiation immunostained for Collagen Type IV (COLIV, green), ZO1 (grey) and F-actin (magenta), showing the morphological process of lumenogenesis and epithelialization. Nuclei were stained with DAPI (blue). Scale bar 25 μm. **E**. Design of the implemented protocol. **F**. Bright field images showing the morphology of differentiating single-cell naïve hiPSCs every 4 days. Scale bar 100 µm. **G**. Principal component analysis (PCA) of bulk RNA-seq of 3D naïve hiPSCs (D0) and 3D naïve hiPSCs at D4, D8, D12, D16 and D20 of neural induction showing the trajectory (grey arrow) of cell identity transition. **H**. Principal component analysis (PCA) of bulk RNA-seq of 3D naïve hiPSCs undergoing neural induction and 3D naïve hiPSCs undergoing priming (dataset from Cesare et al. 2022). **I**. Principal component analysis (PCA) of bulk RNA-seq of 3D naïve hiPSCs undergoing neural induction and primed hPSCs at D0, D7, D12 and D19 of 2D neural induction (dataset from Van de Leemput, Neuron, 2014^43^). **J**. Pearson correlations between naïve hiPSCs during 3D neural induction, conventional primed hiPSCs in standard 2D culture and human embryonic epiblast at different developmental stages, calculated using all expressed genes (schematic drawings of the human embryo were obtained via Biorender) **K**. Heatmap showing the median-centered expression of selected genes, belonging to naïve, naïve and formative, core and primed pluripotency or neuroepithelium categories, ordered by their belonging to the clusters shown in Fig. S1G. The related pie charts represent the percentage of genes that belong to each of cluster 1-5. Only DEGs are presented. Gene lists were compiled from KEGG pathways and published literature^5,15,17,76,77^.

We assessed the differentiation and morphological features of these cysts by immunostaining (**Fig. 1C-D**). Naïve marker KLF17 was downregulated by day (D)4 (**Fig. 1C**), concomitant with lumenogenesis (**Fig. 1D**). Core pluripotency marker OCT4 persisted at D8, whereas by D12 cysts expressed the neuroepithelial marker SOX1 (**Fig. 1C**). These identity transitions were accompanied by organisation of tight junction protein ZO-1 and polarisation of F-ACTIN towards the centre versus deposition of a basement membrane-like layer of Collagen type IV at the outer surface of cysts (**Fig. 1D**), indicating the establishment of proper apico-basal polarity (**Fig. 1D**). Mitotic events were observed at the apical lumen consistent with correct formation of a pseudostratified epithelial tissue architecture (**Fig. 1D D12, S1A**). In total, by D12, 81% of cysts were SOX1+/OCT4-(**Fig. S1B**). Together, this demonstrates that 3D-cultured naïve hiPSCs can promptly and efficiently differentiate into neuroepithelial cysts in a continuum 3D morphogenetic process.

To test the robustness of 3D naïve hiPSC neuroectoderm differentiation, we generated freshly reprogrammed naïve hiPSCs, verified their identity (**Fig. S1C-D**), and confirmed they also underwent coordinated 3D acquisition of neuroepithelial identity and morphology (**Fig. S1E-F**). We further optimised the protocol to extend the differentiation timeline, as would ultimately be required for patient disease modelling. For good viability in longer-term differentiation, we found it helped to extract the cysts from Matrigel at D10 and then culture them in suspension (**Fig. 1E-F**). Subsequently, they formed aggregates with rosette-like internal structures, reminiscent of the organisation of the *in vivo* ventricular zone (**Fig. 1F, D20**), which continued to express SOX1 whilst their morphology was further elaborated (**Fig. S1E-F, D20**).

We investigated cell identity transitions during differentiation through bulk RNA-seq, analysing starting naïve cells and D4, 8, 12, 16 and 20 of differentiation. Principal component analysis (PCA) for all genes showed a clear transcriptional trajectory (**Fig. 1G**), where each timepoint clustered separately and progressively distanced from the naïve control (Naïve 3D) (**Fig. 1G**). We characterised the identity dynamics by comparing this trajectory with that of naïve hiPSCs during 3D priming^22^ (**Fig. 1H**) and of primed hPSCs during standard 2D neural induction^43^ (**Fig. 1I**). We observed alignment of our 3D neural induction trajectory with the priming trajectory for the first 8 days (**Fig. 1H**) and with the primed neural induction trajectory from D4 onward (**Fig. 1I**). This suggests that naïve hiPSCs transiently gain a primed-like signature before transitioning towards the neuroepithelial identity.

*In vivo*, the natural progression of the pre-implantation epiblast is to transition through the post-implantation stage before commencing lineage commitment^1^. We compared the transcriptional signatures of our cells during 3D neural induction with those of human epiblast cells at different embryonic stages (**Fig. 1J**) and observed that our intermediate “primed-like” population (D4-D8) has the highest similarity with the post-implantation epiblast compared also to conventional hiPSCs (**Fig. 1J**).

To further investigate the transition from naïve via transient-primed to neuroepithelial identity in 3D, we analysed differentially expressed genes (DEGs) for all pairwise comparisons between any two timepoints of the naïve-to-neuro trajectory and performed clustering analysis of their dynamics (**Fig. S1G**). We distinguished five major clusters, identified by minimising total variations within each cluster: 1, fast down (904); 2, slow down (3768); 3, up flat (1017); 4, slow up (3468); 5, fast up (993) (**Fig. S1G**). Gene pathway enrichment analysis showed that pluripotency-related categories were among the rapidly downregulated DEGs while neural-related categories were rapidly upregulated (**Fig. S1H**). Consistent with the IF data (**Fig 1C, S1E**), *Sox1* transcript was upregulated from D12 onwards (**Fig. 1K** heatmap). For each cell identity state (naïve pluripotentcy, primed pluripotnecy, formative pluripotency, core pluripotency and neuroepithelium) we analyzed the fraction of genes belonging to the five major cluster of gene dynamics (**Fig. 1K, S1G**). Most of the genes related to naïve pluripotency (100%), shared between naïve and formative pluripotency (100%) and related to core pluripotency (85.7%) are downregulated (slowDN and fastDN clusters). Genes related to primed pluripotency mostly fit into the upregulated clusters (84.6%, UPflat, fastUP and slowUP clusters) but some (15.4%) are downregulated. Importantly, genes related to neuroepithelial categories become upregulated (100%) after 12 days of differentiation (**Fig. 1K**) coherent with the identity dynamics described above (**Fig. 1C-J**).

During development, metabolic rewiring of stem and precursor cells is crucial to regulate their maintenance and differentiation^44^. We observed initial downregulation of genes related to oxidative phosphorylation, coherent with the naïve-to-primed transition (**Fig. S1I**). From D12 genes belonging to both metabolic categories seem to be further downregulated, although the expression of glycolytic genes remained slightly higher (**Fig. S1I**), consistently with the metabolic state reported for neural progenitors^44^. Moreover, analysis of polarity and adhesion-related DEGs showed expression dynamics that support gradual epithelialization and lumenogenesis, associated with the initial primed-like fate acquisition (**Fig. S1J**). Moreover, downregulation of E-cadherin and upregulation of N-cadherin and claudins (**Fig. S1J**), support the emergence of the neuroepithelial identity as these genes *in vivo* play important roles in the morphogenesis of the neural tube^45,46^.

Together these results show that single naïve hiPSCs can differentiate into 3D neuroepithelial cysts in a continuum morphogenetic process that, interestingly, transitions through a primed-like state. This transient state correlates closely with the *in vivo* post-implantation epiblast, more so than conventional/stable primed hPSCs, suggesting that performing neural induction starting from the naïve state might allow more faithful recapitulation of *in vivo* developmental transitions.

### Naïve-derived neuroepithelial cysts acquire specific regional identity along the AP axis

*In vivo* the developing neural tube is exposed to temporal and spatial gradients of morphogens that determine the identity pattern of emerging neural progenitors along the anterior-posterior (AP) axis^47^. To characterise the AP identity of naïve-derived 3D neuroepithelial cysts, we assembled a list of genes spanning four regions of the *in vivo* developing neural tube (Forebrain, Midbrain, Hindbrain, and Spinal cord). We plotted their expression levels in a heatmap (**Fig. 2A**) and observed that almost all were expressed and upregulated during differentiation (**Fig. 2A**). This suggests that in the “default” trajectory, naïve-derived neuroepithelial cells do not uniformly acquire one specific regional identity (**Fig. 2A**).

**Figure 2.**
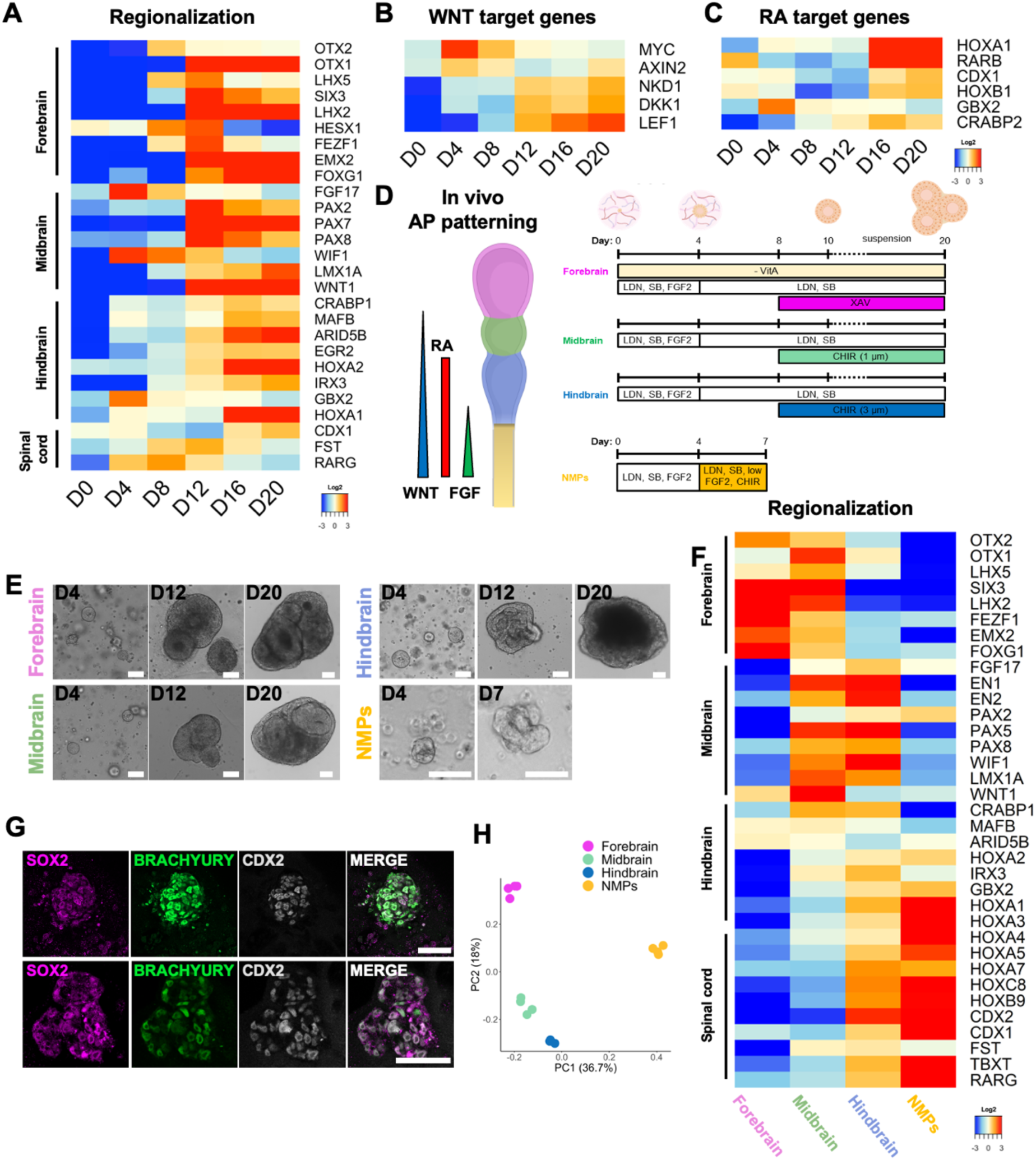
Naïve derived neuroepithelial cysts can acquire specific regional identity along the AP axis. **A**. Heatmaps showing the median-centred expression of selected genes related to the first emerging A-P regions of the developing neural tube (Forebrain, Midbrain, Hindbrain, Spinal cord), for D0, D4, D8, D12, D16 and D20 of 3D neural differentiation. Gene lists were compiled from published literature^77,78^. **B-C**. Heatmaps showing the median-centred expression of WNT βcatenin pathway-(B) or Retinoic acid pathway-(C) target genes. Only DEGs are presented in heatmaps. **D**. Left: schematic representation of human neural tube patterning along the anterior-posterior (A-P) axis into Forebrain (magenta), Midbrain (green), Hindbrain (blue) and Spinal cord (yellow) regions and the morphogens gradients contributing to the patterning. Right: specific *in vitro* differentiation conditions for each brain region or neuromesodermal progenitors (NMPs) are illustrated. RA, Retinoic Acid; -vitA, B27 supplement minus vitamin A; LDN, LDN-193189; SB, SB-431542; FGF2, fibroblast growth factor 2; XAV, XAV939; CHIR, Chiron.). **E**. Bright field images showing the morphology of D4, D12 and D20 of 3D neural differentiating naïve hiPSCs in Forebrain, Midbrain and Hindbrain protocols, and of D4 and D7 of 3D NMP protocol. Scale bar 100 µm. **F**. Heatmaps showing the median-centred expression of selected genes related to the first emerging A-P regions of the developing neural tube at D20 (Forebrain, Midbrain, Hindbrain) or D7 (NMPs) of 3D neural differentiation for the regionalization protocols described in panel D. Gene lists were compiled from published literature^77,78^. **G**. Representative single confocal slices of 3D NMPs at D7 obtained from 3D-cultured naïve hiPSCs following the protocol described in panel D. Colonies were immunostained for SOX2 (magenta), BRACHYURY (green) and CDX2 (grey). Nuclei were stained with DAPI (blue). 2 different cell lines are represented: the BJ naïve line (upper panel) and the HPD03 naïve line (lower panel). Scale bar 50 µm. **H**. PCA of bulk RNA-seq of 3D naïve-derived D20 Forebrain, Midbrain, Hindbrain and D7 NMPs.

AP patterning is mainly controlled by WNT/βcatenin and Retinoic Acid (RA) pathways which specify posterior progenitors within the growing neural tube^47^. We examined the expression dynamics of target genes related to both pathways during 3D neural induction (**Fig. 2B-C**). We observed the upregulation of both WNT and RA target genes during differentiation (**Fig. 2B-C**), suggesting pathway activation. This might explain the mixed AP regional identity signatures of naïve-derived neuroepithelial cysts (**Fig. 2A**). Based on these observations, we designed new differentiation protocols in which we modulated WNT and RA activity to differentiate cells into neuroepithelial progenitors of specific AP identities (**Fig. 2D**). To generate forebrain-like cysts, we removed vitamin A, the precursor of RA, from the basal medium and included XAV-939^48–50^, a WNT inhibitor, from D8 of differentiation (**Fig. 2D**, forebrain), since we observed that WNT targets become upregulated at D12 (**Fig. 2B**). To generate midbrain and hindbrain progenitors, we supplemented the basal medium with a low dose (1 μM, midbrain) or a high dose (3 μM, hindbrain) of CHIR99021 (CHIR), a WNT activator, from D8 (**Fig. 2D**, midbrain and hindbrain). Applying signal modulations from D8 also coincides with the timing of transition from primed-like epiblast to neuroepithelial state, which occurs between D8 and D12 (**Fig 1C, 1H-K, 2A**), suggesting it is the appropriate timing for directing regional identity within the forming neuroepithelium.

Cells exposed to the “forebrain” differentiation protocol first produced cysts with a spherical morphology, a single central lumen and thick epithelium, then generated aggregates with internal rosettes (**Fig. 2E**, forebrain). Cells exposed to “midbrain” and “hindbrain” differentiation conditions grew similarly until CHIR was supplemented to the medium. Then, after 12 days of culture, we observed the formation of structures with multiple elongated lumens and a thinner epithelium (**Fig. 2E**, midbrain and hindbrain). To verify the regional identities, we performed bulk RNA-seq analysis at D20 (forebrain, midbrain and hindbrain) (**Fig. 2F**). We evaluated the expression levels of region-specific genes and plotted them in a heatmap (**Fig. 2F**). We observed distinguishable transcriptional patterns: the forebrain protocol upregulated genes related to forebrain identity (e.g. FOXG1, OTX2 and FEZF1) while the midbrain, hindbrain protocols downregulate forebrain-specific genes and upregulate more posterior genes as appropriate (**Fig. 2F**).

To achieve CNS fates more posterior than the hindbrain, it is necessary to posteriorize the epiblast and induce neuromesodermal progenitors (NMPs), prior to allocation to the spinal cord lineage^3,51^. To test whether 3D-cultured naïve hiPSCs could be differentiated to NMPs, we adapted published protocols for NMP differentiation from primed hPSCs^52,53^, adding the posteriorizing CHIR pulse whilst still in the epiblast continuum. Therefore, we added CHIR on D4 (**Fig. 2D**, NMPs), when cysts most closely represent the PostEPI (**Fig. 1J**). By D7, naïve cells in NMP conditions developed into 3D colonies with compact and apolar morphology (**Fig. 2E**, NMPs), and co-expressed CDX2, BRACHYURY and SOX2 consistent with proper induction of NMP identity (**Fig. 2G**). Bulk RNA-seq analysis of D7 NMPs and comparison with the transcriptomic signatures of forebrain, midbrain and hindbrain by PCA analysis (**Fig. 2H**) showed transcriptional diversity between the samples generated with the different protocols. NMP differentiation resulted in downregulation of forebrain- and midbrain-specific genes and upregulation of more posterior genes (**Fig. 2F**).

WNT and RA target genes were also differentially expressed among cells derived from the different protocols (upregulated only in hindbrain and NMP differentiation conditions) (**Fig. S2A-B**). This suggests that we successfully modulated the pathway activities during the differentiation process thus directing the acquisition of regional identities. Together these results demonstrate that 3D-cultured naïve hiPSCs have the potency to differentiate into neuroepithelial progenitors of different AP identities, upon timely stimulation with morphogenetic signals.

### Naïve-derived neuroepithelial cysts can be matured into forebrain organoids

To test whether naïve hiPSCs can generate brain organoids, we moved forebrain-regionalised (D20) neuroepithelial cysts for further maturation in shaking and normoxic conditions to support nutrient and oxygen exchange (**Fig. 3A**). At D30, we changed the media composition to support neurogenesis and axon sprouting (removal of inhibitors, addition of NT3 and BDNF) (**Fig. 3A**). From D30-60, this led to an increase in neurons (MAP2+ cells, **Fig. 3B**) around the neurogenic niches. The anterior identity of these organoids was confirmed by FOXG1 expression, especially in neural progenitors, and CTIP2 in the surrounding neurons (**Fig. 3C**), a marker of both cortical and striatal forebrain neurons^54,55^.

**Figure 3.**
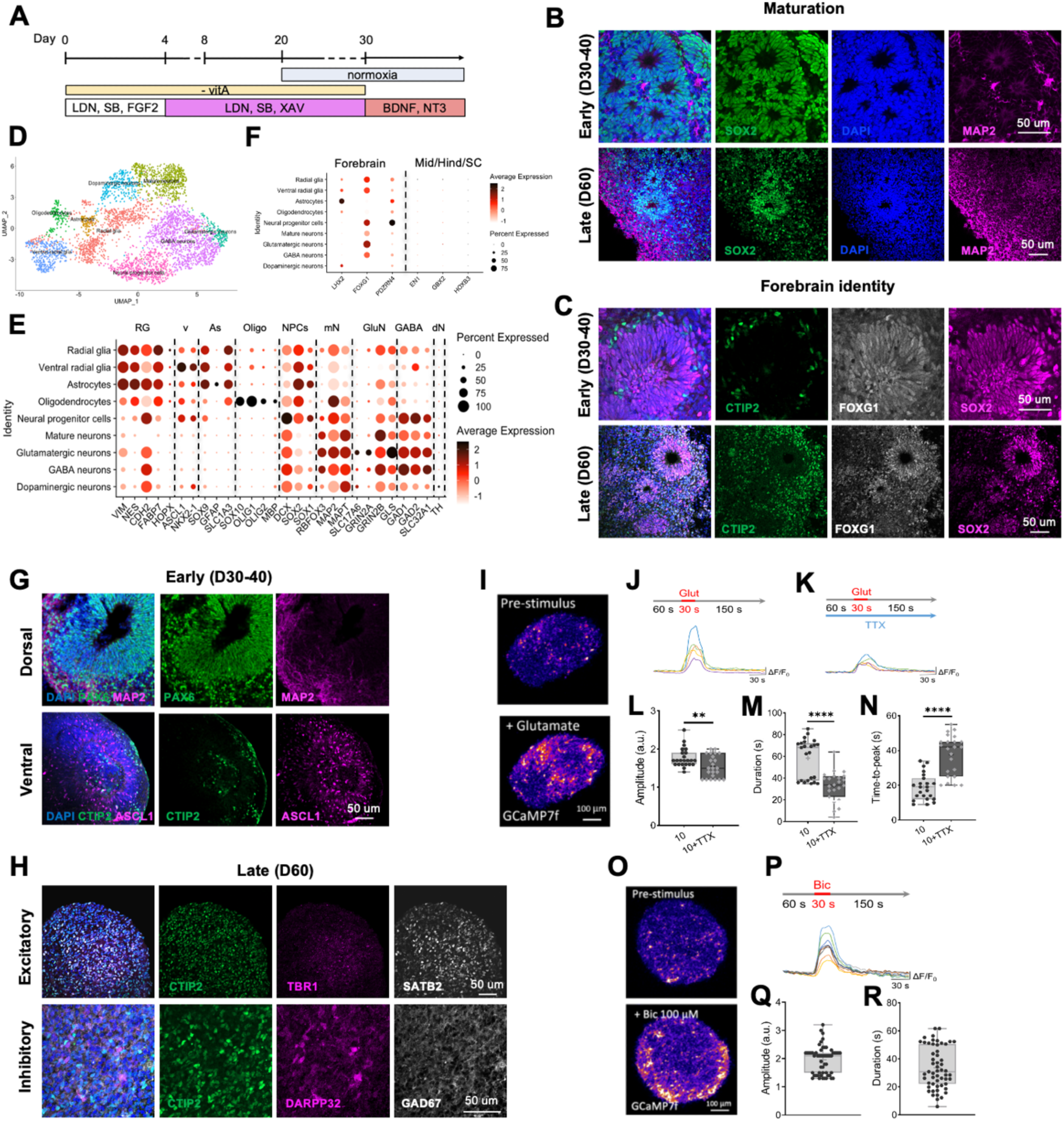
Naïve-derived forebrain neuroepithelial cysts can mature into forebrain organoids including ventral and dorsal identities. **A.** Differentiation protocol conditions to pattern human naïve iPSCs into forebrain organoids. BDNF, Brain-derived neurotrophic factor; NT3, Neurotrophin-3. **B.** Representative confocal images of forebrain organoids at early (D30-45, upper row) and late (D60, lower row) time points stained for SOX2 (neural stem cells, green), DAPI (nuclei, blue) and MAP2 (mature neurons, purple). Scale bars =50 um. **C.** Representative confocal images of forebrain organoids at early (D30-45, upper row) and late (D60, lower row) time points stained for CTIP2 (forebrain neurons, green), FOXG1 (forebrain, grey), SOX2 (neural stem cells, purple). Scale bars =50 um. **D.** UMAP visualization of single-cell RNA expression in forebrain organoids at D60 (D60) of differentiation. **E.** Dot plot of expression pattern of markers of different forebrain cell populations RG= radial glia; v= ventral; As= astrocytes; Oligo= oligodendrocytes; NPCs= neural progenitor cells; mN= mature neurons; GluN= glutamatergic neurons; GABA N= GABAergic neurons; dN= dopaminergic neurons. **F.** Dot plot of expression pattern of region identity markers (Forebrain vs midbrain/hindbrain/spinal cord) expressed in the different cell populations identified by cluster analysis. **G.** Representative confocal images of forebrain organoids at early (D30-40) time points stained for the dorsal neural stem cell/radial glial marker PAX6 (green), mature neuronal marker MAP2 (purple) (upper row), and for ventral neural stem cell/radial glial marker ASCL1 (purple) and forebrain neurons CTIP2 (green). Scale bars =50 um. **H.** Representative confocal images of forebrain organoids at late (D60) time point stained for excitatory neuronal markers (CTIP2, green; TBR1, white; SATB2, grey; upper row) and inhibitory neuronal markers (CTIP2, green; DARP32, purple; GAD67, grey, lower row). Scale bars =50 um. **I.** Representative two-photon images of GCaMP7-GFP infected human forebrain organoids before and after exposure to 10 μM Glutamate. **J-K.** Calcium transients in single neurons upon stimulation for 30 seconds with 10 μM Glutamate (J) or stimulation with glutamate in the presence of TTX (K). **L-N.** Quantification of amplitude (L), duration (M) and time-to-peak (N) of glutamate-evoked calcium events without and with TTX treatment. **O.** Representative two-photon images of GCaMP7-GFP infected human forebrain organoids before and after exposure to 100 μM Bicuculline. **P.** Calcium transients in single neurons upon stimulation for 30 seconds with 100 μM Bicuculline. **Q-R.** Quantification of amplitude (Q) and duration (R) of bicuculline-evoked calcium events.

To analyse cell type composition, we performed single-cell RNA-seq on three D60 forebrain organoids (**Fig. 3D**). Unbiased clustering and annotation using scType method^56^ revealed 9 distinct populations spanning neuronal and glial lineages (**Fig. 3D**), confirmed by dot plot analysis of cell type-specific marker gene expression (**Fig. 3E**): neural progenitors (NPCs), GABAergic neurons (GABA), Glutamatergic neurons (GluN), Dopaminergic neurons (dN), Mature neurons (mN), Radial glial cells (RG), ventral RG (v) (ASCL1, NKX2.2), oligodendrocytes (Oligo) (OLIG1/2, SOX10, MBP), and astrocytes (As) (SOX9, GFAP, SLC1A3). Regional marker expression confirmed these organoids represent forebrain (LHX, FOXG1, PDZRN4) and not midbrain (EN1), hindbrain (GBX2) or spinal cord (HOXB3) (**Fig. 3F**).

We confirmed the presence of both dorso-cortical and ventro-striatal components in naïve-derived forebrain organoids via immunostainings at different timepoints. At D30-40, neurogenic niches expressed the ventral marker ASCL1 (**Fig. 3G**) or dorsal marker PAX6 (**Fig. 3G**). At D60, GABAergic interneurons were marked by GAD67/CTIP2, some of which co-expressed DARP32 indicative of striatal medium spiny neurons (**Fig. 3H**). Different subtypes of glutamatergic excitatory neurons were present expressing TBR1, CTIP2 or SATB2, normally present in layer I-VI, V and II/III respectively in the cerebral cortex (**Fig. 3H**).

To assess the physiological properties of naïve-derived forebrain organoids, we infected them with a GCaMP7f-expressing lentiviral vector to detect neuronal activity. Calcium imaging revealed glutamatergic receptor activity, indicated by calcium transients following local stimulation with 10 μM glutamate (**Fig. 3I-J**). To investigate whether forebrain organoids established a neuronal network, we blocked action potential-dependent communication between neurons using tetrodotoxin (TTX), a Na^+^ voltage-gated channel blocker (**Fig. 3K-N**). This caused a significant reduction in both the amplitude (**Fig. 3L**) and duration (**Fig. 3M**) of glutamate-evoked calcium events, and an increase in the time-to-peak (**Fig. 3N**), suggesting that neuronal activity in organoids in response to glutamate stimulation was not only dependent on the stimulation itself but also on signal propagation between neurons. To test whether NMDA or AMPA glutamate receptor subtypes underpin this synaptic activity, we pharmacologically inhibited them with D-APV or NBQX respectively (**Fig. S3A-D**). Glutamate-evoked calcium transients were significantly reduced in amplitude by both inhibitors (**Fig. S3E-F**). This reduction was more pronounced with D-APV (**Fig. S3G**), suggesting that synaptic activity was mainly mediated by NMDA receptors. Finally, to corroborate the presence of inhibitory neurons shown by immunostaining (**Fig. 3H**), we stimulated forebrain organoids with 100 µM bicuculline to inhibit GABA_A_ receptors (**Fig. 3O-P**). This prompted calcium transients (**Fig. 3O-R**), indicating the presence of a functional inhibitory compartment in these forebrain organoids, since blocking their inhibitory drive triggered increased activity in their target neurons.

Together, these data confirm the ability of naïve-derived organoids to recapitulate key features of forebrain development including the generation of appropriate cellular diversity and tissue-level physiology at D60.

### Naïve-derived forebrain organoids can model the time-regulated epigenetic silencing of *Fmr1* occurring in FXS

To model the timing of epigenetic silencing of *Fmr1* locus in FXS patients we used our naïve-derived forebrain organoid approach (**Fig. 3**). First, we derived isogenic primed and naïve hiPSCs via mRNA-based reprogramming^17,57^ from fully methylated FXS patient fibroblasts with over 270 CGG repeats (**Fig. S4A-B, Fig. 4A**). Whilst in healthy control (CTR) hiPSCs *Fmr1* was expressed in both naïve and primed states (**Fig. 4B**), the acquisition of naïve identity was indispensable to allow FXS cells to transcribe *Fmr1* mRNA (**Fig. 4B**). Correspondingly, naïve but not isogenic primed FXS hiPSCs produced FMRP (**Fig. 4C-D, S4C-D**), albeit at lower protein level than CTR (**Fig. 4C**). Methylation and CGG repeat length analysis of the *Fmr1* locus confirmed that the FXS patient fibroblasts and derivative primed hiPSCs were fully methylated. In contrast, FXS naïve hiPSCs were completely demethylated, while maintaining the CGG amplification length in the range of full mutation (**Fig. 4A**). Therefore, our naïve FXS hiPSC model captures the hallmarks of *Fmr1* locus epigenetic regulation (**Fig. 4A-B**) and FMRP production (**Fig. 4C-D**). Culturing demethylated FXS naïve iPSCs in proliferative conditions maintains a stable CGG repeat number up to passage (P)10; longer expansion induced a strong contraction in the CGG repeat number (**Fig. 4A**). Therefore, only naïve FXS iPSCs under P10 were used as the starting material.

**Figure 4.**
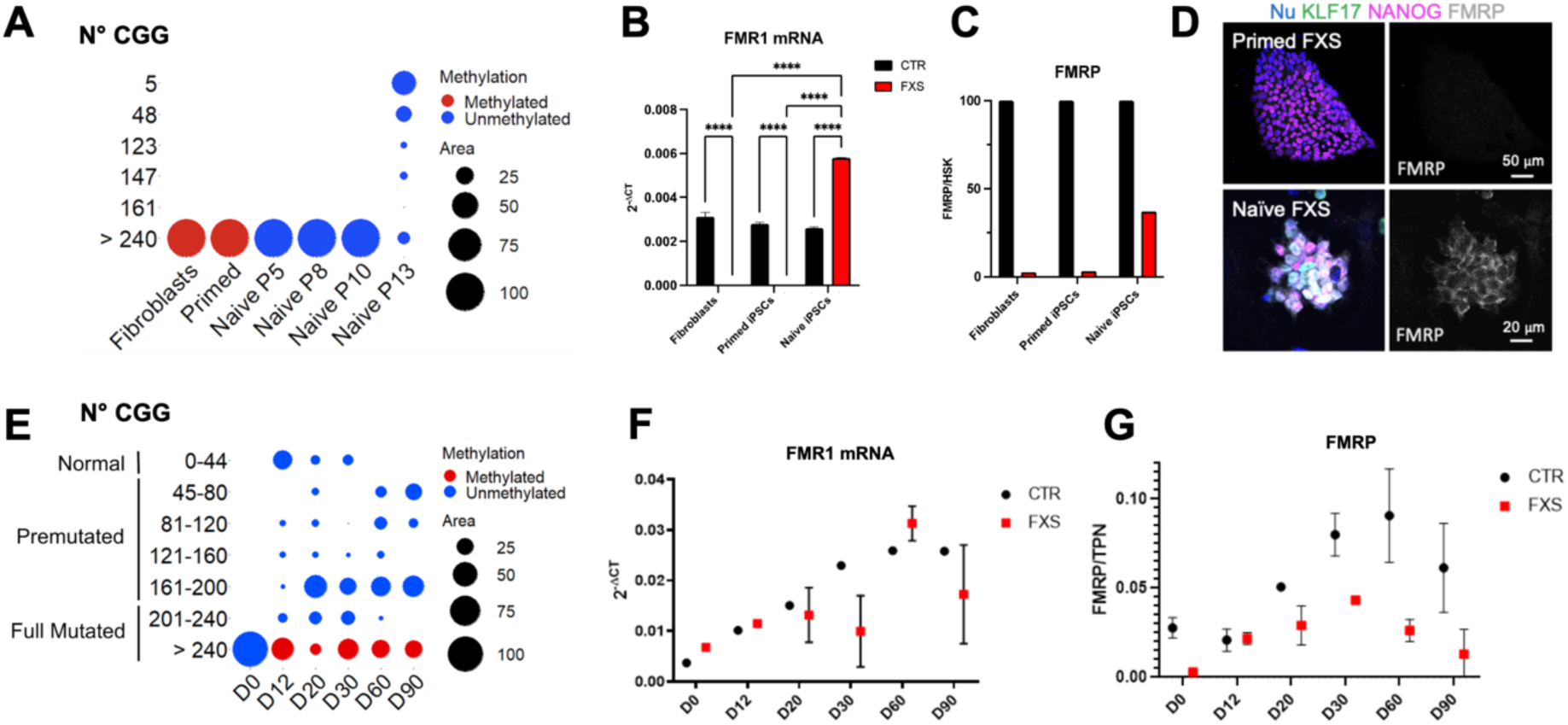
Naïve derived brain organoids can model the early stages of FXS brain development. **A.** Dotplot showing the amplification length and methylation status of CGG repeats at *Fmr1* locus in isogenic fibroblasts, primed and naïve hiPSCs at different passages. Blue are unmethylated alleles, red are methylated. **B.** qRT-PCR for *Fmr1* gene expression in naïve and primed iPSCs from healthy control (CTR, black) and FXS patient (FXS, red) normalised to *Gapdh.* **C.** Western Blot analysis for FMRP protein on fibroblasts, primed and naïve iPSCs from control (CTR, black) and FXS patient (FXS, red). Shown are the values normalised to housekeeping (HSK) protein Actin. **D.** Confocal images of primed (upper) and naïve (lower) hiPSCs from FXS patient. Shown are pluripotency marker NANOG (magenta), the naïve specific marker KLF17 (green) and FMRP (grey). Nuclei were stained with DAPI (blue). **E.** Dotplot representing all the allele sizes generated during differentiation of naïve iPSCs into brain organoids and their methylation status over time (from D0 (naïve iPSCs) to D90). Blue are unmethylated alleles, red are methylated alleles. **F.** qRT-PCR for *Fmr1* gene expression on brain organoids at different time points of differentiation (from D0 to D90) from CTR (black) and FXS patient (red) normalised to *Gapdh.* A pool of minimum 5-6 organoids were used per time point. **G.** Western Blot analysis for FMRP protein on brain organoids at different time points of differentiation (from D0 to D90) from CTR (black) and FXS patient (red). Shown are the values normalised to Ponceau Staining as total protein normalization (TPN). A pool of minimum 5-6 organoids were used per time point.

Having confirmed the complete removal of the epigenetic barrier in our FXS naïve hiPSCs, we differentiated them to forebrain organoids and analysed their methylation patterns and CGG expansion over time. Starting from fully mutated demethylated naïve FXS cells, during differentiation we observed the appearance of alleles of different lengths covering the entire mutational spectrum of the *Fmr1* locus, as observed in naïve iPSCs upon prolonged culture (**Fig. 4A**). However, in forebrain organoids, only alleles with >240 CGG repeats were preferentially methylated (**Fig. 4E**), whereas alleles in naïve cells were always demethylated irrespective of the amplification length of the CGG repeats (**Fig. 4A**). This methylation was observed already at the neuroectodermal stage (D12) (**Fig. 4E**) consistent with the disappearance of FMRP in some neuroectodermal cysts (**Fig. S4E**). These genetic and epigenetic patterns resulted in a specific *Fmr1* transcriptional signature and FMRP synthesis pattern in FXS samples. Indeed, CTR forebrain organoids showed a progressive increase over time in *Fmr1* mRNA transcription and FMRP synthesis, reaching a plateau around D30 of differentiation (**Fig. 4F-G**). In FXS forebrain organoids, *Fmr1* mRNA showed similar behaviour to CTR samples until D12. After D12, the dynamic of *Fmr1* showed higher variability, which correlates with the emergence of pre-mutated alleles, known to produce higher levels of *Fmr1* mRNA compared to normal alleles^26–28^ (**Fig. 4F**). Conversely, FMRP level was lower since the beginning, increased with a slower dynamic until D30, and then decreased almost back to starting levels (**Fig. 4G, S4F**). This trend may result from *Fmr1* transcripts derived from fully mutated unmethylated, and premutated alleles, which may be inefficiently translated into FMRP. Together, these data show that FXS naïve iPSC-derived brain organoids reveal potential pathogenic events associated with genomic instability and the complex dynamic of *Fmr1* silencing and FMRP production during the early phase of brain development.

## Discussion

In this study, we show that 3D-cultured naïve hiPSCs can differentiate into regionalised brain organoids, which can serve as a model for the epigenetically regulated neurodevelopmental disease FXS. We demonstrate that applying neural induction signals to single naïve hiPSCs cultured in a 3D ECM-rich environment induces efficient differentiation, following a developmental timing coherent with *in vivo* human embryogenesis and achieving neuroepithelial identity within 12 days from the naïve state. This is coupled with morphological progression from apolar naïve cells to epithelialisation and apical lumenogenesis then acquisition of pseudostratified neuroepithelial tissue architecture in a continuum 3D morphogenetic process. In parallel, cells transition through sequential stages of pluripotency before neuroepithelial identity acquisition, consistent with the need of naïve cells to acquire competence for germ layer induction via capacitation^15^ and with the transition dynamics of *in vivo* epiblast development^1^. Between D4 and D8, our 3D differentiation better mimics the *in vivo* post-implantation epiblast compared to stable expanded primed hPSCs, as also observed in capacitated cells^58^. Therefore, our approach allows us to generate lineage-specific models, following an *in vivo*-like trajectory and overcoming some epigenetic limitations of primed hPSCs. Furthermore, by achieving germ layer differentiation directly from naïve hiPSCs in 3D, we observed accompanying dynamics related to morphological features and metabolism, including polarity and adhesion-related genes supporting initial epithelialisation and lumenogenesis, then claudins and N-cadherin important for *in vivo* neuroepithelial integrity, apico-basal polarity and neural tube closure^45,46^.

Given the potency of human naïve PSCs to generate both embryonic and extraembryonic tissues and the absence of defined protocols to direct naïve hiPSCs toward 3D neuroectoderm, here we developed a strategy to induce neural fate from 3D naïve hiPSCs by adapting the dual-SMADi regime commonly used for primed hiPSCs. SB431542 (TGFβ inhibitor) treatment promotes 2D-cultured naïve hiPSCs to acquire trophectoderm (TE) fate^59^. Here we found that, when applied from the beginning in combination with LDN (BMP inhibitor) as part of the pro-neuroectodermal dual-SMADi regime^48,49^ we obtain 3D naïve hiPSC differentiation without overt TE induction. The addition of FGF2 likely supports priming and blocks TE, since inhibition of ERK/MAPK also promotes TE induction from naïve hiPSCs^60^. The inclusion of at least one passage in RSeT medium before differentiation (see Material & Methods) may also facilitate embryonic rather than extraembryonic differentiation since it supports a naïve phenotype closer to the peri-implantation epiblast^61^. Furthermore, Matrigel was recently shown to promote epiblast identity in mouse ICM explants due to “inside” positional cue from laminin-integrin α6β1 signalling^62^. In the future, it will be important to dissect how signalling cues and ECM environments contribute individually and in collaboration to elicit the development of a 3D neuroepithelium, including whether local entrapment of paracrine signals and mechanical cues play a role. In human brain organoids, extrinsic matrix provision improves tissue morphogenesis^63^ including timely and proper spatial maturation of cell type complexity^64^. Thus, our 3D model provides a platform to investigate whether the ECM network regulates cellular specification and tissue patterning throughout development from pre-implantation to CNS formation.

Starting from naïve human iPSCs, the epiblast counterpart before any axial patterning, we could induce differentiation of neuroepithelial cysts with distinct axial identities by timely signal modulation (**Fig. 2**). WNT inhibition or titrated activation steered differentiation to forebrain, mid/hindbrain, or NMPs, according to the role of WNT in axis posteriorisation *in vivo*^51,65^. Further maturation from anterior neuroepithelial cysts allowed us to create forebrain organoids from naïve hiPSCs. By D60, these comprised appropriate cell-type complexity including radial glial cells, astrocytes, oligodendrocytes, neural progenitor cells and mature neurons^66^. We found both excitatory and inhibitory neurons, the two main components of mammalian cerebral cortex^67^, which are crucial for proper neural circuit formation^68^ and derive from distinct neurogenic niches^69^ observed in our D60 organoids (**Fig 3**). Thus, 3D naïve hiPSCs can model the timely development of the complex human forebrain.

The hypomethylated state of naïve hiPSCs allowed us to model the genetic and epigenetic dynamics at the *Fmr1* locus during the early stages of FXS development. We reprogrammed primary fibroblasts from FXS patient into naïve hiPSCs and achieved a complete erasure of *Fmr1* epigenetic memory, as previously reported^40,41^, while preserving the CGG expansion within the full-mutated range (**Fig 4A**)^39,40^. Interestingly, transcriptional reactivation of the *Fmr1* locus in FXS naïve hiPSCs led, upon prolonged culture, to CGG repeat contraction and repeat length mosaicism. This phenomenon may be linked to R-loop formation and recruitment of endogenous DNA repair machinery, which results in the shortening of the CGG repeat expansion, as previously demonstrated^40^.

In FXS patients, pathogenic silencing of *Fmr1* is thought to occur during the late first/early second trimester of human embryonic development^30–34,70^. By differentiating FXS patient naïve hiPSCs versus CTR naïve hiPSCs to forebrain organoids, extended to D90 to cover the first trimester, we found that, in FXS naïve-derived forebrain organoids, *Fmr1* methylation was restricted to full-mutated alleles (>240 CGG repeats). This methylation occurred very early during differentiation, being observable as early as D12 in neuroepithelial cells, and possibly even earlier (**Fig. 4E**). This aligns with the observation that FXS primed hESCs exhibit variable methylation^38,39^ and highlights the importance of conducting studies in the peri-implantation phase. Concomitant with the methylation of full-length alleles, we observed CGG repeat contraction and repeat length mosaicism among unmethylated alleles (**Fig. 4A**), further supporting the idea that methylation of full-mutated alleles may stabilise expanded repeats^31,39^.

With naïve hiPCs-derived organoids, we can follow *Fmr1* expression and FMRP synthesis dynamics starting from *Fmr1* expressing naïve cells, as it occurs during embryogenesis, contrary to previously reported models generated from primed hiPSCs that are hypermethylated and never express *Fmr1*^36,37^. *Fmr1* mRNA is widely transcribed and translated into FMRP throughout embryonic brain development and is important in neurogenesis ^71,72^. We found a progressive increase at both mRNA and protein levels in CTR organoids during time, but not FXS organoids, in which *Fmr1* mRNA and FMRP levels reflected averages of cells harbouring different CGG expansions and methylation patterns (**Fig. 4E-G**). Full-mutated and methylated alleles produce neither *Fmr1* mRNA nor FMRP, whereas pre-mutated alleles are known to have an increased *Fmr1* mRNA transcription to counterbalance inefficient FMRP synthesis^26–28^. This mechanism could also be present in full-mutated unmethylated alleles where the full-length mRNA is translated inefficiently, resulting in mRNA accumulation but reduced synthesis of FMRP. This could explain why, in our FXS naïve hiPSC-derived brain organoids containing all these mutational spectra, FMRP levels are reduced throughout differentiation compared to CTR, while *Fmr1* mRNA shows greater variability (**Fig. 4F-G**).

The advantages of differentiating brain organoids from naïve hiPSCs extend beyond their epigenetic *tabula rasa* and differentiation trajectory comprising peri-implantation stages. This approach opens the possibility of creating clonal organoids from single patient cells, which is particularly important for genetic conditions presenting mosaicism. The clonal nature of naïve hiPSCs represents a breakthrough for brain organoid-based disease models, allowing the generation of organoids with specific genetic and epigenetic profiles for each mutational spectrum. This allows the dissection of the contribution of full-mutated and pre-mutated alleles in the early pathogenesis of FXS, and the investigation of when and how mosaicism arises during development. In future studies, naïve hiPCS-derived FXS forebrain organoids will serve as a powerful model to investigate the mechanisms of FXS pathogenesis and disease physiology on the cellular and tissue levels. These insights will be essential for designing new therapeutic strategies to improve the treatment of this complex disease.

## Acknowledgements

We are grateful to Teresa Rayon for advice regarding NMP differentiation protocols, to Irene Zorzan and Graziano Martello for advice regarding the culture of naïve hiPSCs, to Keisuke Ishihara regarding 3D neurectoderm differentiation and to Andrea Ballabio for carefully regarding the manuscript. We thank the support of TIGEM NGS and Bioinformatics Cores and Next Generation Diagnostic srl. C.L. was supported by Marie Curie Individual fellowship (839753) and from FRAXA Foundation Fellowship. H.T.S. was supported by a Wellcome Trust Sir Henry Wellcome postdoctoral fellowship, and by the Austrian Science Fund (FWF) Special Research Area Program (10.55776/F7803-B) to E.M.T.. M.D. was supported by FRAXA Foundation Fellowship. O.G. was supported by the University of Padova under the 2019 STARS Grants program (iNeurons) and Fondazione Umberto Veronesi fellowship. S.B was supported by the MSCA Seal of Excellence @ UNIPD 2023 Postdoctoral Fellowship (project number C95F21010280001) and by Walter Benjamin Postdoctoral Fellowship from the Deutsche Forschungsgemeinschaft (DFG, German Research Foundation) (Project number 524585667). This work was supported by Fondazione Telethon and the European Research Council Advanced Grant (grant agreement 759154, Reproids). J.B. is supported by the Francis Crick Institute, funded by Cancer Research UK, the UK Medical Research Council and the Wellcome Trust (all under CC001051). For the purpose of Open Access, the authors have applied a CC BY public copyright license to any Author Accepted Manuscript (AAM) version arising from this submission.

## Author contributions

C.L., E.C., H.T.S., and N.E. designed the experiments and wrote the manuscript. N.E. supervised the project. C.L., E.C., H.T.S. and M. D. performed and analysed the experiments. A.G. and M.G.L.B. performed RNA-seq bioinformatic analyses. C.R. helped in the characterisation of FMRP dynamic in hiPSCs and organoids. A.Maset performed and analysed calcium imaging experiments. S.B. helped in long-term differentiation of naïve-derived brain organoids. O.G., S.A. and R.F. optimised naive hiPSC reprogramming in microfluidics and helped analyse the data. J.B. and E.M.T. advised during creation of 3D neurectoderm differentiation protocols from naïve hiPSCs and during manuscript preparation. R.P. performed sizing and methylation analysis of FMR1 locus and helped in data interpretation. A.Murgia helped in FXS data interpretation and contributed to manuscript preparation. D.C. performed RNA-sequencing and helped in data interpretation.

## Methods

### Cell lines

Human naïve induced pluripotent stem cells (naïve hiPSCs, HPD06, HPD03 and FXS GM05131)^17^, primed induced pluripotent stem cells (hiPSCs; HPD06 and FXS GM05131) and human fibroblasts (HFF-1 and BJ from ATCC, and GM05131 from Coriell Institute Biobank) were used for this study. The transcriptomic, metabolic and epigenetic profiles of HPD06 and HPD03 naïve lines were extensively characterised in Giulitti et al.^17^ to confirm their similarity to other naïve cell lines. The human primed HPD06 line was generated and characterised in Cesare et al.^22^. Human naïve and primed iPSC lines from HFF, BJ and FXS patients were specifically generated and characterised within this study.

### Cell reprogramming in microfluidics

Naïve and primed hiPSCs from FXS, HFF and BJ fibroblasts were generated by modifying the protocol previously described^17^. Microfluidic chips are prepared in-house according to the procedure already described^57,73^.

### Cell culture in 2D

Pluripotent cell lines were cultured at 37°C, 5% CO2 and 5% O_2_ atmosphere while fibroblasts were maintained at 37°C and 5% CO_2_ atmosphere. Cell lines were routinely tested and confirmed negative for mycoplasma. Naïve hiPSCs were cultured on a confluent layer of mitotically inactive MEFs (DR4, ATCC) in PXGL medium^74^. Cells were split every 4-5 days with TrypLE Select Enzyme (Life Technologies) as previously described. Primed hiPSCs were expanded in wells (Corning) coated with 0.5% Matrigel growth factor reduced (MRF, Corning), cultured in Essential 8 medium (E8, Stemcell Technologies) and passaged every 4-5 days with EDTA (ThermoFisher Scientific). Media change was performed every 24 hours (h) for both naïve and primed hiPSCs. Human fibroblasts were cultured on 100 mm Petri dishes (Falcon) in High Glucose Dulbecco’s modified Eagle Medium (DMEM, ThermoFisher Scientific) supplemented with 10% Fetal Bovine Serum (FBS, ThermoFisher Scientific). Medium was changed every second day.

### Cell expansion in 3D

Naïve cells were switched from 2D to 3D culture by detaching naïve colonies from 2D MEF, dissociating the colonies as single cells with TrypLE and then embedding them in 50% Matrigel (Corning) 50% RSeT (Voden) drops at 2k cells/µl. For maintenance, 25 µl of the Matrigel/cell mixture was spread on tissue culture treated multiwell plates (Falcon) and jellified at 37°C. RSeT medium was used to feed the embedded cells, including Rock inhibitor (Ri) (10 μM) (Myltenyi) for the first day. Medium change was performed every 24 h and cells were split after 4-5 days for direct 3D differentiation. To split 3D cultured cells, we follow the procedure previously described^22^.

### Naïve to organoid conversion in 3D

To induce naïve iPSCs toward a neural organoid identity in 3D, we developed the following protocol. Naïve cells previously cultured in 3D Matrigel, in RSeT medium for 1 passage were dissociated to single cells with TrypLE and resuspended in 100% Matrigel at 2000 cells/μl. 10 μl of cell suspension was then seeded as a thin drop, jellified, and fed with advanced N2B27 medium to promote robust neural organoid differentiation and growth^75^. Medium comprises Advanced DMEMF12 (Life Technologies), Neurobasal (Life Technologies), B27 (Life Technologies) (1:100), N2 (Life Technologies) (1:200), L-Glutamine (Life Technologies) (1:100), non essential amino acids (NEAA, Life Technologies) (1:100), β-mercaptoethanol (bMe, Life Technologies) (1:1000), PenStrep (Life Technologies) (1:100). We further supplemented advanced N2B27 with FGF2 (20 ng/mL, PeproTech), LDN-193189 (LDN, Sigma) (10-100 nM), SB-431542 (SB, Axon Medchem) (10 μM) and Ri (10 μM). After 48h Ri was withdrawn and, 5 days after seeding, bFGF was also withdrawn such that cultures proceeded in advanced N2B27+LDN+SB. To obtain anterior forebrain identities we applied the same medium compositions but using B27 without vitamin A (Life Technologies) and supplementing with XAV939 (Axon Medchem, 5 µM) from D8 of culture. To obtain more posterior brain regions CHIR (Abcr GmbH) was supplemented from D8 of culture at two different concentrations (1 µM or 3 µM). Medium was changed daily. At D10 organoids were removed from Matrigel and placed in cell culture suspension dishes, while at D20 they were moved to an orbital shaker (Thermo Fisher), set at 70 r.m.p., at 37°C, 5% CO2 and 21% O2 atmosphere. From D30, advanced N2B27 medium was supplemented with BDNF (20 ng/µl, Preprotech) and NT3 (20 ng/µl, Preprotech) and 1% Matrigel, whilst LDN and SB were withdrawn. To generate neuromesodermal progenitors (NMPs), cells were treated for the first 4 days of culture as described above for non AP patterned organoids. From D4 to D7, medium was replaced with advanced N2B27 medium + LDN + SB + FGF2 (5 ng/ml) + CHIR (3 µM). On D7 cultures were stopped for analyses.

### Immunofluorescence

Naïve, primed and fibroblast cells were cultured on glass coverslips (12 mm^2^) and fixed 4 days after seeding for immunofluorescence characterization. Naïve cells were seeded on MEF coated coverslips and primed cells or fibroblasts on coverslips coated with 2.5% MRF. Cells were fixed with 4% paraformaldehyde (PFA) (Sigma-Aldrich) in PBS without calcium and magnesium (-/-), 10 min at room temperature. Brain organoids still embedded (untill D10) were fixed in 2% PFA for 45 minutes, whereas brain organoids derived from suspension culture were fixed in 4% PFA for 1h at room temperature. All samples were then washed 3 times with PBS -/- and stored at 4°C in PBS+P/S until used.

For immunostaining of 2D samples, permeabilization and blocking were performed simultaneously by incubating samples with 5% Normal Donkey Serum (Sigma-Aldrich) in PBS -/- with 0.1% (v/v) Triton X-100 (Sigma-Aldrich) for 1 h at room temperature. Then cells were incubated overnight at 4°C with primary antibodies, diluted in blocking solution. The day after, samples were washed with PBS -/- three times for 5 min and then incubated with secondary antibodies, diluted in blocking solution. Incubation was performed for 1h at room temperature. Samples were washed 3 times with PBS-/- and then mounted with Fluoromount-G (Southern Biotech). 3D cultured cells and organoids were treated with 5% normal donkey serum (Sigma-Aldrich) in PBS -/- with 0.5% (v/v) Triton X-100 for 3 h at room temperature for permeabilization and blocking. Then samples were incubated 24 h at 4°C with primary antibodies, diluted in blocking solution. The day after, samples were washed with PBS -/- with 0,5% Triton three times for 15 min and then incubated 24 h at 4°C with secondary antibodies, diluted in blocking solution. Samples were then washed three times with PBS -/- with 0,5% Triton for 15 min, then once with PBS -/- for 15 min. Samples were adapted to and then mounted in Fluoromount-G (Southern Biotech).

All primary and secondary antibodies used are listed in **Table 1**. Nuclei were stained with DAPI (Thermo Scientific) and F-ACTIN with Phalloidin 488 or 647 (Invitrogen). Confocal fluorescence images were acquired at a LSM900 confocal microscope (Zeiss).

To prepare histology samples, after fixation organoids were cryoprotected in 30% Sucrose (Sigma-Aldrich) 24h at 4°C and embedded in OCT (Sakura Finetek) for snap-freezing. Samples were then cut at cryostat in 20 μm-thick slices. For immunostaining, slices were washed 3 times in PBS-/-to remove OCT. At this point, depending on the primary antibody combination, some samples underwent antigen retrieval step, so they were incubated in Dako Target Retrieval Solution, Citrate pH 6 1X (Agilent) at 70°C for 40 minutes, they were washed three times in PBS -/- and then permeabilised and blocked simultaneously by incubating samples with 5% normal donkey serum in PBS with 0.1% (v/v) Triton X-100 for 1h at room temperature. Then, primary antibodies diluted in blocking solution were incubated overnight at 4°C. The day after, samples were washed three times for 5 min in PBS with 0.1% (v/v) Triton X-100 and then incubated with secondary antibodies diluted in blocking solution for 1h at room temperature. Before mounting, samples were washed three times in PBS and then mounted using DABCO (homemade solution made of glycerol, Polyvinyl alcohol, and DABCO, Sigma-Aldrich).

### Viral infection

To allow GCaMP7f expression in organoids for multi-photon functional imaging, organoids were infected with pLV[Exp]-Puro-SYN1>jGCaMP7f generated by VectorBuilder. 2 weeks before the imaging organoids were transferred to 1.5 ml Eppendorf tubes. Next, 15 µl of culture medium containing the lentiviral vector (1 × 10^8^ TU/ml titration) in the presence of polybrene (1 µg/ml) was added to the tube, and the organoids were moved to the incubator for 60 min. Next, 85 µl of medium was added to each tube, and the tubes were returned to the incubator overnight. The next day, organoids were washed with fresh medium and transferred to low-attachment plates. The medium was changed two times per week.

### Multi-photon functional imaging

Organoids’ neuronal activity recordings were acquired using a two-photon microscope (Scientifica 2P, Uckfield, United Kingdom), equipped with a Ti:Sapphire laser (Chameleon Ultra II, Coherent) light source tuned at 920 nm to excite the genetically encoded calcium indicator GCaMP7f. Images were acquired at 1.72 frames per second, 512×512 pixel resolution, with a water dipping 16x objective (Nikon LWD DIC N2, N.A. 0.8). Fluorescence emission was filtered by a main dichroic mirror (525 −556 nm / 580 −650 nm, Olympus) in combination with a 525/39 nm short pass emission filter (Olympus) and collected by a photomultiplier tube (GaAsP, Scientifica). For the characterisation of the neuronal activity, organoids were placed into an imaging chamber (RC-26, Warner Instruments) and continuously perfused with oxygenated (5% CO_2_) Ringer solution (NaCl 140 mM, KCl 5mM, CaCl_2_ 1 mM, MgCl_2_ 1 mM, HEPES 10 mM, Na-Pyruvate 1 mM, Glucose 10 mM, penicillin/streptomycin 1% v/v, pH 7.3) at 37°C with a flow rate of 1 ml/min. The stimulation protocol consisted of a 60-s baseline followed by a 30-s pulse of 100 µl of glutamate 10 µM (Tocris), and a 150-s washout, for a total imaging time of 4 min. To pharmacologically modulate the network activity the following inhibitors were added to Ringer solution, incubated for 15 min before the imaging session, and continuously perfused with the Ringer solution throughout the entire imaging session: Tetrodotoxin (TTX 5 µM, Tocris), D-(-)-2-Amino-5-phosphonopentanoic acid (D-APV 50 µM, Tocris), 2,3-Dioxo-6-nitro-1,2,3,4-tetrahydrobenzo[*f*]quinoxaline-7-sulfonamide (NBQX 20 µM, Tocris). After the imaging session, the inhibitor was washed with fresh Ringer solution three times by completely changing the medium in the imaging chamber and subsequently leaving the organoid under continuous perfusion with fresh Ringer solution for 15 min. The sequence of D-APV and NBQX incubation was randomic.

### Functional imaging analysis and quantification

Time series with functional activity were processed with ImageJ software for motion correction and manual segmentation of active cells. For each cell, the extracted fluorescence trace was processed by subtracting the background noise and normalizing the fluorescence trace on the median intensity value of the first 30 frames. To identify and quantify glutamate-evoked calcium transients, data were binarized based on a threshold defined as the median intensity value + 2.5 standard deviations of the first minute of fluorescence trace. The amplitude, duration, starting time, and the time-to-peak of the calcium transients were calculated for each cell. To assess the effect of TTX application, the mean amplitude, duration, and time-to-peak of the glutamate-evoked activity for each organoid before and after TTX application were compared using the Wilcoxon test. The cumulative distribution of the starting time of each cell was fitted to a sigmoidal distribution. The goodness of fit was assessed by calculating the coefficient of determination (R^2^), and the probability for the data belonging to the non-TTX and TTX datasets to fit the same sigmoidal distribution was calculated. To assess the effect of D-APV and NBQX application, the mean amplitude of the glutamate-evoked activity for each organoid before and after D-APV and NBQX application was compared using the Wilcoxon test. The mean relative amplitude reductions upon D-APV and NBQX application for each organoid were then compared using the Mann-Whitney test. All the datasets were analysed with Prism 8 software (GraphPad).

### Bulk RNA sequencing and transcriptomic analysis

RNA sequencing data shown in **Fig. 1** and **Fig. S1** were obtained from 3D naïve hPSCs and 3D naïve during neural differentiation in 3D Matrigel at D4, 8, 12, 16 and 20 of differentiation. RNA sequencing data shown in **Fig. 2** were obtained on region-specific neural cysts at D20 (Forebrain, Midbrain and Hindbrain) and D7 (NMPs). Four replicates were collected for each sample. To collect the RNA samples, naïve colonies and 3D differentiating cells were extracted from the Matrigel drops then lysed with RLT buffer. RNA was extracted using the RNeasy Plus Mini Kit (Qiagen). Total RNA was quantified using the Qubit 2.0 fluorimetric Assay (Thermo Fisher Scientific).

RNA-sequencing of these samples was performed by the Next Generation Sequencing Facility of the Telethon Institute of Genetics and Medicine (TIGEM) in Napoli. Libraries were prepared from 600-1000 pg of total RNA using the SMART-Seq v4 Ultra Low Input Kit (Takara Bio) followed by Nextera XT DNA Library Preparation kit (Illumina). Libraries were sequenced on a NovaSeq 6000 sequencing system using a paired-end, 150-cycle strategy (Illumina Inc.) for the samples of the neural differentiation time course and using a single-end, 100-cycles strategy (Illumina Inc.) for samples of the region-specific neural cysts. Illumina Illumina NovaSeq 6000 base call (BCL) files were converted in fastq file through bcl2fastq. Data obtained from the sequencing were analysed in our lab at the University of Padova. Alignment was performed with STAR 2.6.0a124 using hg38 GENECODE reference genome and the expression levels of genes were determined with RSEM 1.3.0. Genes were annotated with the Ensembl database and raw counts were expressed as counts per million (CPM). For the samples of the neural differentiation time course, data was filtered to keep only genes with more than 10/min (library size*10^-6) count per million (CPM) in more than three replicates in at least one sample. For samples of the region-specific neural cysts, all the genes that did not have at least 3 CPM in more than three replicates in at least one sample were filtered out. Data was normalised by negative binomial distribution (TMM) with EdgeR (v 3.40.2). Differentially expressed genes (DEGs) analysis was performed using EdgeR on processed data, using a mixed criterion based on p value, after false discovery rate (FDR) correction by Benjamini Hochberg method, lower than 0.05 and absolute log2(fold change) higher than 2. Integration of the literature dataset was performed on log2(CPM+1) data with the ComBat-seq tool using the sva (v 3.46.0) R package. PCAs in **Fig. 1G, H, I** and **Fig. 2H** were performed on log2(CPM+1) data using R (v. 4.1.3). PCA in **Fig. S1C** was performed on log2(CPM+1) data using MATLAB R2019b (The MathWorks).

The union of all found DEGs in every paired combination of samples from the neural differentiation time course was used for further analyses. DEGs were assigned to five clusters, as in Rostovskaya et al., Dev. 2019^15^, using k-means algorithm implemented in MATLAB R2019b (The MathWorks) and the gene’s temporal profiles for each of the clusters were represented.

For the 3D neural differentiation samples, enrichment analyses within GO-BP, KEGG and Reactome databases were performed using the ClueGo plug-in of Cytoscape (v 3.9.1) with Benjamini Hochberg corrected p-value lower than 0.05. Enrichment results were presented in waterfall plots (**Fig. S1H**) highlighting categories related to pluripotency and neural identities. The lists of the categories, per each plot, are reported in **Table 2**. Pearson’s correlation coefficients were calculated between 3D neural differentiation data and pseudobulk gene expression of the embryonic epiblast subpopulations obtained from Sousa et al., 2023^58^. Heat map data visualization was performed using the median-centered expression of a list of selected genes based on published literature.

### Single-cell RNA sequencing and transcriptomic analysis

Data shown in **Fig. 3** were obtained from organoids at D60 of maturation. Three organoids were mechanically minced in small pieces and dissociated at single cell by incubating with Papain and DNase I solution (Papain Dissociation System, Worthington Biochemical) for 40 minutes at 37°C, with shaking in separated plates. Digestion was blocked using the Ovomucoid protease inhibitor (Papain Dissociation System, Worthington Biochemical) and cells were pelleted by centrifugation. Cell pellet was resuspended in PBS-/-supplemented with a Running buffer containing BSA 0.4% and sodium azide (Milteny) and filtered through a cell strainer to remove cell clumps and debris. Cells were counted and diluted to the desired density of 1000 cells/µl. 20000 cells for each of the three organoids were pooled together for the subsequent analysis.

Libraries were prepared following the 10X Single Cell 3’ v2 protocol. Sequencing was performed on the Illumina Novaseq 6000 platform at the Next Generation Sequencing Facility of the Telethon Institute of Genetics and Medicine (TIGEM) in Naples. Data were analysed in our lab, at the University of Padova. Pre-processing of data is performed with Cellranger software. Cellranger pipeline together with 10X standard Chromium barcode sequences are used for the generation of fastq files. The same pipeline is used for alignment, filtering, barcode, and counting of unique molecular identifiers (UMI), and alignment to human genome reference (GRCh38/hg38 reference genomes). A dataset of 27001 cells was obtained with Seurat (v. 4.3.0) standard workflow in RStudio (v 2022-10-31). Cells having less than 1000 and more than 7500 detected genes, read count greater than 10000 and mitochondrial associated reads percentage greater than 5% were filtered out. After this step 3851 good quality cells were obtained. Values of gene expression are normalised to CPM and transformed to the log2 scale using a pseudocount of 1 to standardize the expression counts. Seurat’s ScaleData function was used regressing out the percentage of mitochondrial genes detected.

Principal Component Analysis (PCA) on the scaled data was computed and non-linear dimension reduction using uniform manifold projection (UMAP) was performed using 15 principal components. Graph-based clustering was performed using the FindNeighbors and FindClusters Seurat functions with a resolution of 0.7. Automated cluster annotation was performed according to the ScType method^56^.

### Quantitative reverse transcriptase PCR

Cell lysates were collected from both 2D and 3D-cultured samples by incubating them with RLT buffer (QIAGEN). Total RNA was then extracted using RNeasy mini kit, according to the manufacturer protocol (QIAGEN) including the RNase-free DNase (QIAGEN) step to digest contaminant DNA. cDNA was produced from <300 ng RNA using the SuperScript^TM^ VILO^TM^ cDNA Synthesis kit, following the recommended protocol (Invitrogen). RT-qPCR reactions were performed using the Taqman^TM^ Gene Expression Master Mix (Thermo Fisher Scientific) and Taqman^TM^ Gene Expression assays (Thermo Fisher Scientific, listed in **Table 3**). Three technical replicates were performed for all qPCR experiments. GAPDH was used as the endogenous control to normalise the gene expression.

### CGG repeat length and methylation analysis

Cell lysates were collected from hiPSCs (naïve and primed) and organoids at different time points (D0, 12, 20, 30, 60, 90). DNA was extracted via QIAamp® DNA mini Kit and quantified by spectrophotometer with NanoDrop^TM^ (ThermoFisher Scientific, Waltham, MA, USA). DNA samples were analysed for methylation status and the amplification of the CGG repeats using the AmplideX *Fmr1* mPCR reagents (Asuragen) according to the manufacturer’s recommended protocol. Briefly, 160 ng DNA samples were premixed with two plasmids: a digestion control (DigCtrl) and PCR reference control (RefCtrl). This premix was separately aliquoted to a control or methylation-sensitive digestion reaction. All alleles were detected using FAM-labelled primers, but only the proportion of the protected methylated allele was available for PCR using HEX-labelled primers. Lack of methylation at either HpaII site resulted in digestion and thus no amplification. All amplicons were analysed by capillary electrophoresis (CE) on a 3130xl Genetic Analyzer (Applied Biosystems, ThermoFisher Scientific, Waltham, MA, USA) equipped with POP-7 polymer and a 36 cm array. GeneMapper® v 4.0 software with ROX 1000 size ladder (Asuragen, Austin, TX, USA) was used for automated sizing of *Fmr1* gene-specific peaks in conjunction with mobility correction factors.

### Western Blot

Proteins from 3D-cultured samples at different timepoints of the neural differentiation protocol were extracted using 50 μl of lysis buffer (50mM Tris HCl pH 7.4, 150mM NaCl (Sigma - S3014-1KG), 1% NP-40, 0.5% sodium deoxycholate (Merck – 30970-25G), 0.1% Sodium dodecyl sulphate (Merck - 71736-100ML), 1mM EDTA (Gibco/Thermo – 15575020), 10mM NaF, 1x cOmplete™ EDTA-free Protease Inhibitor Cocktail (Sigma-4693132001)). After sonication (3×30’’ max power) and centrifugation (13000 g 10’ +4°C), the supernatants were quantified using Pierce BCA Protein Assay Kit (Thermo Scientific – 23227)). For western blot analysis, proteins were resolved by 4-12% bis-Tris Gel (NuPAGE) and transferred onto nitrocellulose membrane. Protein transfer was checked by Ponceau S staining (Sigma) and a picture was acquired for further Total Protein Normalization (TPN). The membrane was blocked with 5% non-fat dry milk in TBS-Tween 1% and incubated with FMRP (1:100 Mouse Santa Cruz) and Actin (1:8000 Rabbit Merck) primary antibodies overnight at 4°C. Membrane was washed 3 times and incubated with secondary peroxidase antibodies (1:1000, Bio-RAD) 1 hour in TBS-Tween 1%, followed by washing and chemiluminescence revelation (Immobilon Classico Western HRP substrate - Sigma WBLUC0500). Images were acquired by Alliance Q9 Advance Chemidoc (UVITEC) and analysed on ImageJ.

---

**Supplementary Figure 1.**
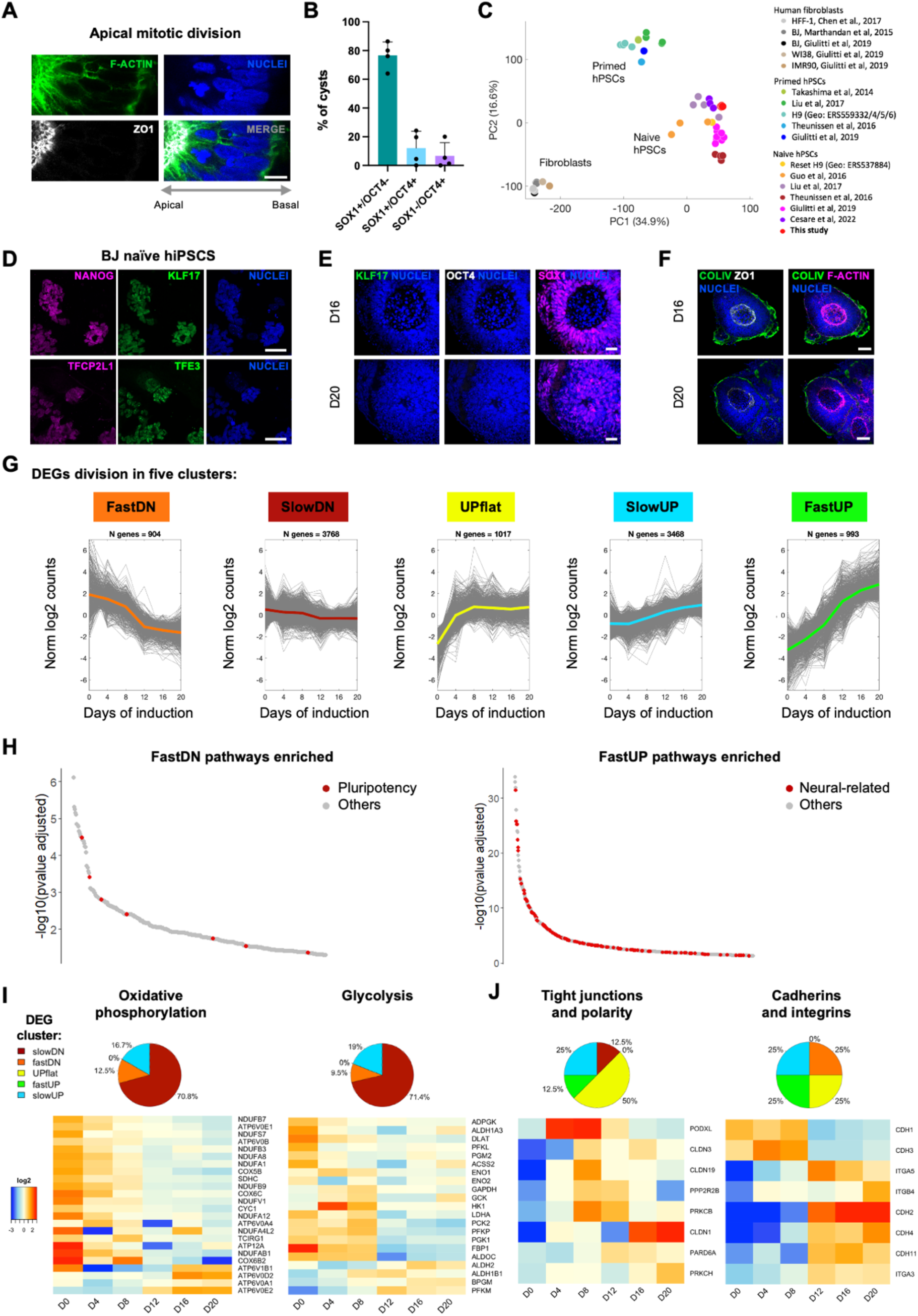
Characterisation of freshly reprogrammed naïve hiPSCs and their expression dynamics during transition to neuroepithelial cysts in 3D. **A**. Representative single confocal slice of D12 3D neural differentiation immunostained for ZO1 (grey) and F-actin (green) showing apical localization of mitotic nuclei. Nuclei were stained with DAPI (blue). Scale bar 10 μm. Zoom from Fig. 1D D12. **B**. Quantification of the percentage of SOX1+/OCT4-, SOX1+/OCT4+ and SOX1-/OCT4+ cysts at D12 of 3D neural differentiation. Plot bars show the mean value + s.d., n = 4. **C**. PCA of newly generated naïve hiPSCs (red) cultured in 3D Matrigel. Human fibroblasts, primed hiPSCs, and naïve hiPSCs from other studies were used as controls. Names of the cell lines are reported in **Table 4**. **D**. Representative single confocal slices of naïve hiPSCs culture immunostained for KLF17 and TFE3 (green), NANOG and TFCP2L1 (magenta). Nuclei were stained with DAPI (blue). Scale bars, 50 mm. **E**. Representative single confocal slices of D16 and D20 3D neural differentiation immunostained for KLF17 (green), OCT4 (grey) and SOX1 (magenta). Nuclei were stained with DAPI (blue). Scale bar 25 μm. **F**. Representative single confocal slices of D16 and D20 3D neural differentiation immunostained for Collagen Type IV (COLIV, green), ZO1 (grey) and F-actin (magenta). Nuclei were stained with DAPI (blue). Scale bar 50 μm. **G**. Temporal expression profiles of DEGs compiled from all pairwise comparisons between any two samples divided by clustering analysis into five dynamic clusters. **H**. Waterfall plots showing the most significant enriched pathways, selected from BP, KEGG and Reactome databases, in the fastDN (left panel) and FastUP (right panel) clusters. Pluripotency-related and neural-related categories are highlighted in the plots (red dots). The categories used to generate the waterfall plots are listed in **Table 2**. **I-J**. Heatmaps showing the median-centred expression of genes related to metabolism (I) and tissue polarity (J) and their belonging to the clusters shown in panel. G. The related pie charts represent the percentage of genes that belong to each cluster. Only DEGs are presented in heatmaps. Gene lists were compiled from KEGG pathways and published literature^15,17,76^.

**Supplementary Figure 2.**
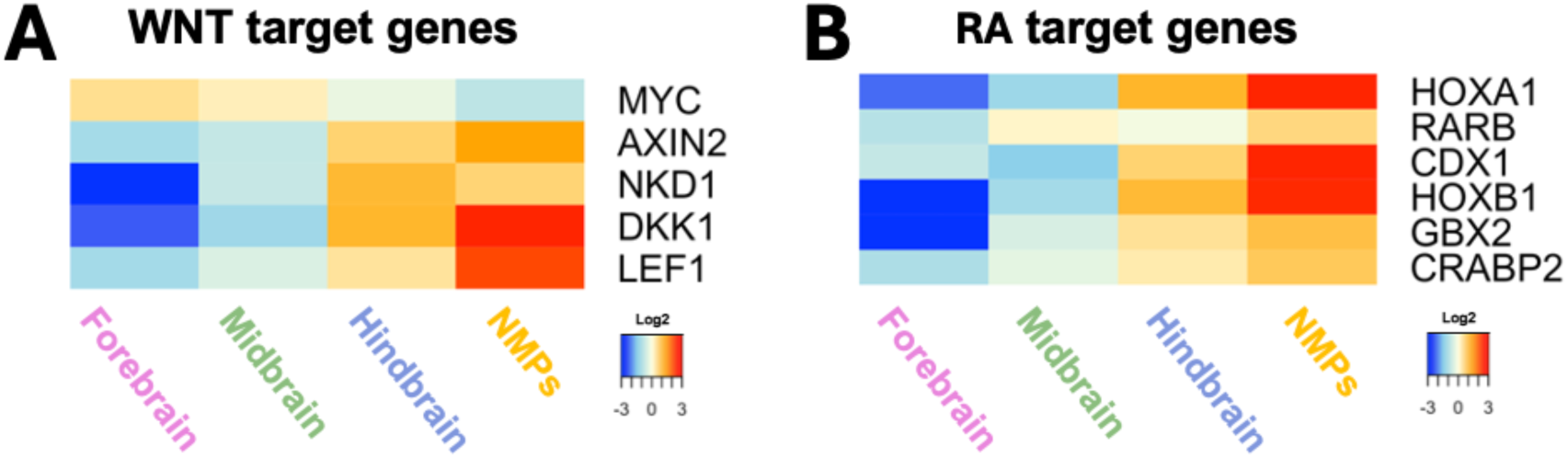
Differential expression of WNT and RA targets in organoids representing the different AP idendities. Heatmaps showing the median-centred expression of WNT βcatenin pathway-(A) or Retinoic acid pathway-(B) target genes in organoids with different AP identities. Only DEGs are presented in heatmaps.

**Supplementary Figure 3.**
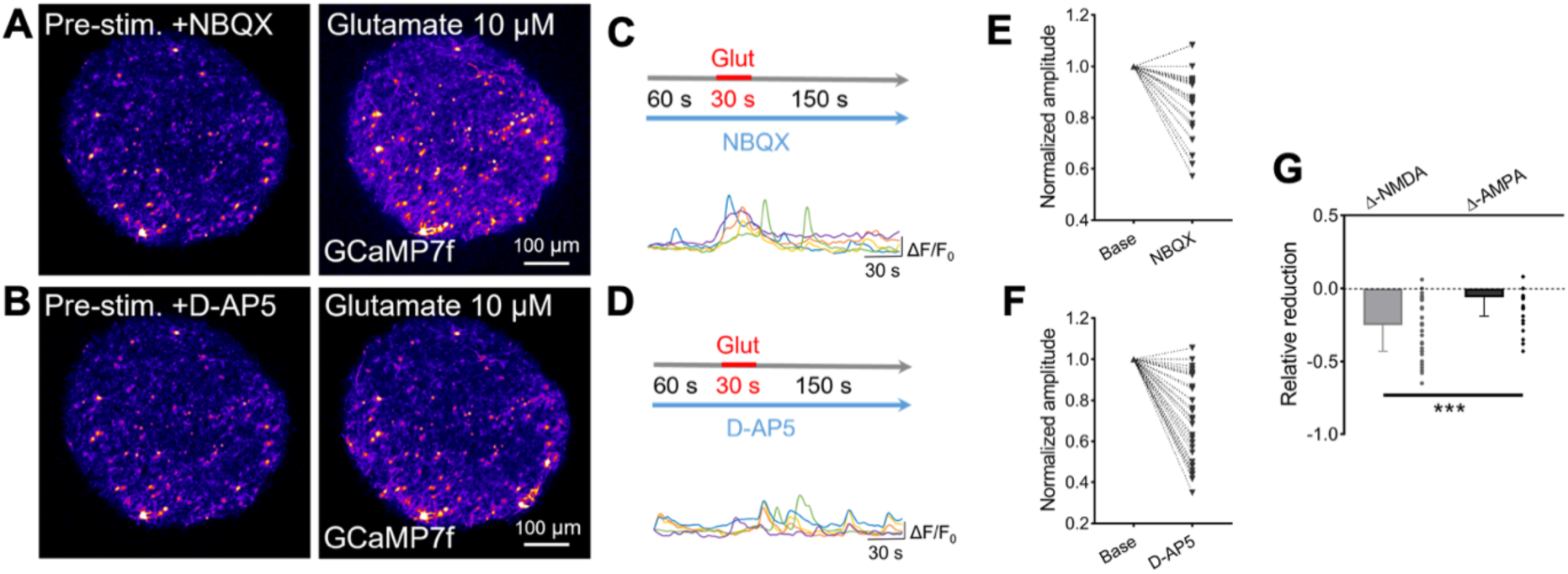
Electrophysiological properties of cortical brain organoids. **A-B**. Representative two-photon stop-frame images of GCaMP7-GFP infected human forebrain organoids before and after exposure to 10 μM glutamate in the presence of AMPA receptor inhibitor NBQX (A) and NMDA receptor inhibitor D-AP5 (B). **C-D**. Calcium transients in single neurons upon stimulation for 30 seconds with 10 μM glutamate in the presence of NBQX (C) or D-AP5 (D). **E-F**. Analysis of amplitude variation upon treatment with NBQX (E) or D-AP5 (F). **G**. Relative contribution of NMDA and AMPA receptors to the reduction in glutamate-evoked neural response.

**Supplementary Figure 4.**
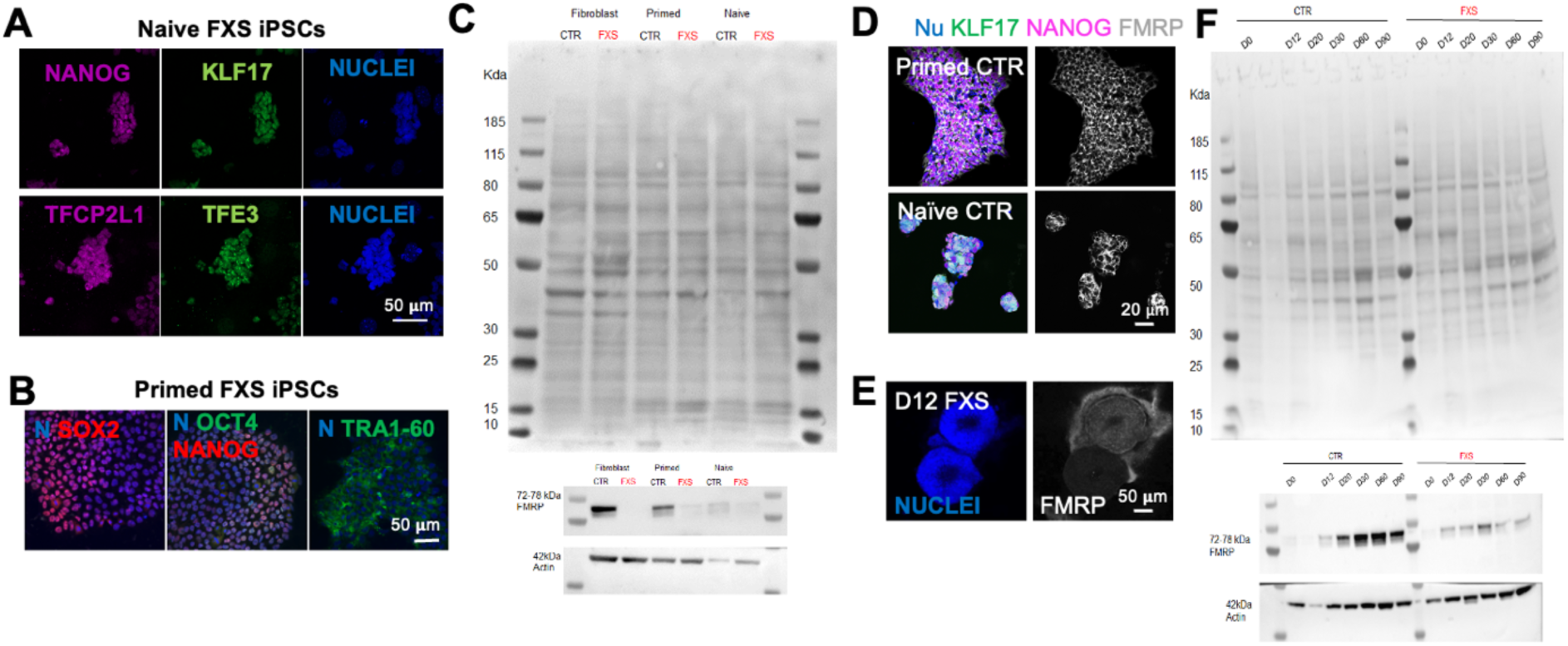
Naïve derived brain organoids can model the early stages of FXS brain development. **A.** Representative single confocal slices of naïve iPSCs after 4 days in culture immunostained for KLF17 and TFE3 (green), NANOG and TFCP2L1 (magenta). Nuclei were stained with DAPI (blue). Scale bars, 50 mm. **B.** Confocal images of primed FXS iPSCs immunostained for SOX2 (red), OCT4 (green) and NANOG (red), TRA1-60 (GREEN). Nuclei are counterstained with DAPI (blue). **C.** Representative Western Blot membrane images for Ponceau Staining, FMRP, ACTIN of FXS and CTR fibroblasts, primed and naïve. **D.** Confocal images of primed (upper) and naïve (lower) iPSCs from healthy control (BJ naïve and primed iPSCs). Shown are pluripotency marker NANOG (magenta), the naïve specific marker KLF17 (green) and FMRP (grey). Nuclei are counterstained with DAPI (blue, N). **E.** Representative image of brain organoid at D12 of differentiation showing heterogeneous presence of FMRP (white). Nuclei are counterstained with DAPI (blue). **F.** Representative Western Blot membrane images for Ponceau staining, FMRP, ACTIN of FXS and CTR organoids at time points D0, 12, 20, 30, 60, 90 of differentiation.

## References

1. Rossant, J., and Tam, P.P.L. (2022). Early human embryonic development: Blastocyst formation to gastrulation. Dev Cell 57, 152–165. 10.1016/J.DEVCEL.2021.12.022.

2. Ozair, M.Z., Kintner, C., and Brivanlou, A.H. (2013). Neural induction and early patterning in vertebrates. Wiley Interdiscip Rev Dev Biol 2, 479–498. 10.1002/WDEV.90.

3. Tzouanacou, E., Wegener, A., Wymeersch, F.J., Wilson, V., and Nicolas, J.F. (2009). Redefining the Progression of Lineage Segregations during Mammalian Embryogenesis by Clonal Analysis. Dev Cell 17, 365–376. 10.1016/j.devcel.2009.08.002.

4. Gouti, M., Metzis, V., and Briscoe, J. (2015). The route to spinal cord cell types: a tale of signals and switches. Trends in Genetics 31, 282–289. 10.1016/J.TIG.2015.03.001.

5. Lancaster, M.A., Renner, M., Martin, C.A., Wenzel, D., Bicknell, L.S., Hurles, M.E., Homfray, T., Penninger, J.M., Jackson, A.P., and Knoblich, J.A. (2013). Cerebral organoids model human brain development and microcephaly. Nature 2013 501:7467 501, 373–379. 10.1038/nature12517.

6. Kelley, K.W., and Pașca, S.P. (2022). Human brain organogenesis: Toward a cellular understanding of development and disease. Cell 185, 42–61. 10.1016/J.CELL.2021.10.003.

7. Osafune, K., Caron, L., Borowiak, M., Martinez, R.J., Fitz-Gerald, C.S., Sato, Y., Cowan, C.A., Chien, K.R., and Melton, D.A. (2008). Marked differences in differentiation propensity among human embryonic stem cell lines. Nature Biotechnology 2008 26:3 26, 313–315. 10.1038/nbt1383.

8. Bock, C., Kiskinis, E., Verstappen, G., Gu, H., Boulting, G., Smith, Z.D., Ziller, M., Croft, G.F., Amoroso, M.W., Oakley, D.H., et al. (2011). Reference maps of human es and ips cell variation enable high-throughput characterization of pluripotent cell lines. Cell 144, 439–452. 10.1016/J.CELL.2010.12.032/ATTACHMENT/4B6CE50B-455E-43C2-92C7-3121A3D64C99/MMC8.PDF.

9. Kim, K., Zhao, R., Doi, A., Ng, K., Unternaehrer, J., Cahan, P., Hongguang, H., Loh, Y.H., Aryee, M.J., Lensch, M.W., et al. (2011). Donor cell type can influence the epigenome and differentiation potential of human induced pluripotent stem cells. Nature Biotechnology 2011 29:12 29, 1117–1119. 10.1038/nbt.2052.

10. Ohi, Y., Qin, H., Hong, C., Blouin, L., Polo, J.M., Guo, T., Qi, Z., Downey, S.L., Manos, P.D., Rossi, D.J., et al. (2011). Incomplete DNA methylation underlies a transcriptional memory of somatic cells in human iPS cells. Nature Cell Biology 2011 13:5 13, 541–549. 10.1038/ncb2239.

11. Sherman, S., Pletcher, B.A., and Driscoll, D.A. (2005). Fragile X syndrome: Diagnostic and carrier testing. Genetics in Medicine 7, 584–587. 10.1097/01.GIM.0000182468.22666.DD.

12. Zhou, J., Hu, J., Wang, Y., and Gao, S. (2023). Induction and application of human naive pluripotency. CellReports 42, 112379. 10.1016/j.celrep.2023.112379.

13. Thorold Theunissen, A.W., Friedli, M., He, Y., Trono, D., Ecker, J.R., and Jaenisch, R. (2016). Molecular Criteria for Defining the Naive Human Pluripotent State Accession Numbers GSE75868. Cell Stem Cell 19, 502–515. 10.1016/j.stem.2016.06.011.

14. Buckberry, S., Liu, X., Poppe, D., Tan, J.P., Sun, G., Chen, J., Nguyen, T.V., de Mendoza, A., Pflueger, J., Frazer, T., et al. (2023). Transient naive reprogramming corrects hiPS cells functionally and epigenetically. Nature 2023 620:7975 620, 863–872. 10.1038/s41586-023-06424-7.

15. Rostovskaya, M., Stirparo, G.G., and Smith, A. (2019). Capacitation of human naïve pluripotent stem cells for multi-lineage differentiation. Development 146. 10.1242/DEV.172916.

16. Guo, G., von Meyenn, F., Rostovskaya, M., Clarke, J., Dietmann, S., Baker, D., Sahakyan, A., Myers, S., Bertone, P., Reik, W., et al. (2017). Epigenetic resetting of human pluripotency. Development (Cambridge) 144, 2748–2763. 10.1242/dev.146811.

17. Giulitti, S., Pellegrini, M., Zorzan, I., Martini, P., Gagliano, O., Mutarelli, M., Ziller, M.J., Cacchiarelli, D., Romualdi, C., Elvassore, N., et al. (2018). Direct generation of human naive induced pluripotent stem cells from somatic cells in microfluidics. Nature Cell Biology 2018 21:2 21, 275–286. 10.1038/s41556-018-0254-5.

18. Warrier, S., Van Der Jeught, M., Duggal, G., Tilleman, L., Sutherland, E., Taelman, J., Popovic, M., Lierman, S., Chuva De Sousa Lopes, S., Van Soom, A., et al. (2017). Direct comparison of distinct naive pluripotent states in human embryonic stem cells. Nat Commun. 10.1038/ncomms15055.

19. Dupont, C. (2024). A comprehensive review: synergizing stem cell and embryonic development knowledge in mouse and human integrated stem cell-based embryo models. Front Cell Dev Biol 12, 1386739. 10.3389/FCELL.2024.1386739/BIBTEX.

20. Pedroza, M., Gassaloglu, S.I., Dias, N., Zhong, L., Hou, T.C.J., Kretzmer, H., Smith, Z.D., and Sozen, B. (2023). Self-patterning of human stem cells into post-implantation lineages. Nature 2023 622:7983 622, 574–583. 10.1038/s41586-023-06354-4.

21. Kagawa, H., Javali, A., Khoei, H.H., Sommer, T.M., Sestini, G., Novatchkova, M., Scholte op Reimer, Y., Castel, G., Bruneau, A., Maenhoudt, N., et al. (2021). Human blastoids model blastocyst development and implantation. Nature 2021 601:7894 601, 600–605. 10.1038/s41586-021-04267-8.

22. Cesare, E., Urciuolo, A., Stuart, H.T., Torchio, E., Gesualdo, A., Laterza, C., Gagliano, O., Martewicz, S., Cui, M., Manfredi, A., et al. (2022). 3D ECM-rich environment sustains the identity of naive human iPSCs. Cell Stem Cell 29, 1703–1717.e7. 10.1016/j.stem.2022.11.011.

23. Hagerman, R.J., Berry-Kravis, E., Hazlett, H.C., Bailey, D.B., Moine, H., Kooy, R.F., Tassone, F., Gantois, I., Sonenberg, N., Mandel, J.L., et al. (2017). Fragile X syndrome. Nature Reviews Disease Primers 2017 3:1 3, 1–19. 10.1038/nrdp.2017.65.

24. Gheldof, N., Tabuchi, T.M., and Dekker, J. (2006). The active FMR1 promoter is associated with a large domain of altered chromatin conformation with embedded local histone modifications. Proc Natl Acad Sci U S A 103, 12463–12468. 10.1073/PNAS.0605343103/SUPPL_FILE/05343FIG6.PDF.

25. Richter, J.D., and Zhao, X. (2021). The molecular biology of FMRP: new insights into fragile X syndrome. Nature Reviews Neuroscience 2021 22:4 22, 209–222. 10.1038/s41583-021-00432-0.

26. Tassone, F., Hagerman, R.J., Taylor, A.K., Mills, J.B., Harris, S.W., Gane, L.W., and Hagerman, P. (2000). Clinical involvement and protein expression in individuals with the FMR1 premutation. Am J Med Genet 91, 144–152. 10.1002/(SICI)1096-8628(20000313)91:2<144::AID-AJMG14>3.0.CO;2-V.

27. Tassone, F., Hagerman, R.J., Taylor, A.K., Gane, L.W., Godfrey, T.E., and Hagerman, P.J. (2000). Elevated Levels of FMR1 mRNA in Carrier Males: A New Mechanism of Involvement in the Fragile-X Syndrome. The American Journal of Human Genetics 66, 6–15. 10.1086/302720.

28. Kenneson, A., Zhang, F., Hagedorn, C.H., and Warren, S.T. (2001). Reduced FMRP and increased FMR1 transcription is proportionally associated with CGG repeat number in intermediate-length and premutation carriers. Hum Mol Genet 10, 1449–1454. 10.1093/HMG/10.14.1449.

29. Pretto, D., Yrigollen, C.M., Tang, H.T., Williamson, J., Espinal, G., Iwahashi, C.K., Durbin-Johnson, B., Hagerman, R.J., Hagerman, P.J., and Tassone, F. (2014). Clinical and molecular implications of mosaicism in FMR1 full mutations. Front Genet 5, 103594. 10.3389/FGENE.2014.00318/ABSTRACT.

30. Castellví-Bel, S., Milà, M., Soler, A., Carrió, A., Sánchez, A., Villa, M., Jiménez, M.D., and Estivill, X. (1995). Prenatal diagnosis of fragile x syndrome: (cgg)n expansion and methylation of chorionic villus samples. Prenat Diagn 15, 801–807. 10.1002/PD.1970150903.

31. Devys, D., Biancalana, V., Rousseau, F., Boue, J., Mandel, J.L., and Oberle, I. (1992). Analysis of full fragile X mutations in fetal tissues and monozygotic twins indicate that abnormal methylation and somatic heterogeneity are established early in development. Am J Med Genet 43, 208–216. 10.1002/AJMG.1320430134.

32. Willemsen, R., Bontekoe, C.J.M., Severijnen, L.A., and Oostra, B.A. (2002). Timing of the absence of FMR1 expression in full mutation chorionic villi. Hum Genet 110, 601–605. 10.1007/S00439-002-0723-5/METRICS.

33. Suzumori, K., Yamauchi, M., Seki, N., Kondo, I., and Hori, T.A. (1993). Prenatal diagnosis of a hypermethylated full fragile X mutation in chorionic villi of a male fetus. J Med Genet 30, 785–787. 10.1136/JMG.30.9.785.

34. Sutcliffe, J.S., Nelson, D.L., Zhang, F., Pieretti, M., Caskey, C.T., Saxe, D., and Warren, S.T. (1992). DNA methylation represses FMR-1 transcription in fragile X syndrome. Hum Mol Genet 1, 397–400. 10.1093/HMG/1.6.397.

35. Colvin, S., Lea, N., Zhang, Q., Wienisch, M., Kaiser, T., Aida, T., and Feng, G. (2022). 341 Repeats Is Not Enough for Methylation in a New Fragile X Mouse Model. eNeuro 9. 10.1523/ENEURO.0142-22.2022.

36. Kang, Y., Zhou, Y., Li, Y., Han, Y., Xu, J., Niu, W., Li, Z., Liu, S., Feng, H., Huang, W., et al. (2021). A human forebrain organoid model of fragile X syndrome exhibits altered neurogenesis and highlights new treatment strategies. Nature Neuroscience 2021 24:10 24, 1377–1391. 10.1038/s41593-021-00913-6.

37. Raj, N., McEachin, Z.T., Harousseau, W., Zhou, Y., Zhang, F., Merritt-Garza, M.E., Taliaferro, J.M., Kalinowska, M., Marro, S.G., Hales, C.M., et al. (2021). Cell-type-specific profiling of human cellular models of fragile X syndrome reveal PI3K-dependent defects in translation and neurogenesis. Cell Rep 35. 10.1016/J.CELREP.2021.108991/ATTACHMENT/02B6DF91-61A2-4692-921D-F9ADBE4AE244/MMC2.PDF.

38. Avitzour, M., Mor-Shaked, H., Yanovsky-Dagan, S., Aharoni, S., Altarescu, G., Renbaum, P., Eldar-Geva, T., Schonberger, O., Levy-Lahad, E., Epsztejn-Litman, S., et al. (2014). FMR1 epigenetic silencing commonly occurs in undifferentiated fragile X-affected embryonic stem cells. Stem Cell Reports 3, 699–706. 10.1016/j.stemcr.2014.09.001.

39. Eiges, R., Urbach, A., Malcov, M., Frumkin, T., Schwartz, T., Amit, A., Yaron, Y., Eden, A., Yanuka, O., Benvenisty, N., et al. (2007). Developmental Study of Fragile X Syndrome Using Human Embryonic Stem Cells Derived from Preimplantation Genetically Diagnosed Embryos. Cell Stem Cell 1, 568–577. 10.1016/j.stem.2007.09.001.

40. Lee, H.G., Imaichi, S., Kraeutler, E., Aguilar, R., Lee, Y.W., Sheridan, S.D., and Lee, J.T. (2023). Site-specific R-loops induce CGG repeat contraction and fragile X gene reactivation. Cell 186, 2593–2609.e18. 10.1016/J.CELL.2023.04.035/ASSET/704D972C-8B75-45CF-B62D-786A44E2074F/MAIN.ASSETS/GR7_LRG.JPG.

41. Gafni, O., Weinberger, L., Alfatah Mansour, A., Manor, Y.S., Chomsky, E., Ben-Yosef, D., Kalma, Y., Viukov, S., Maza, I., Zviran, A., et al. (2013). Derivation of novel human ground state naive pluripotent stem cells. Nature 504. 10.1038/nature12745.

42. Chambers, S.M., Fasano, C.A., Papapetrou, E.P., Tomishima, M., Sadelain, M., and Studer, L. (2009). Highly efficient neural conversion of human ES and iPS cells by dual inhibition of SMAD signaling. Nature Biotechnology 2009 27:3 27, 275–280. 10.1038/nbt.1529.

43. van de Leemput, J., Boles, N.C., Kiehl, T.R., Corneo, B., Lederman, P., Menon, V., Lee, C., Martinez, R.A., Levi, B.P., Thompson, C.L., et al. (2014). CORTECON: A temporal transcriptome analysis of in vitro human cerebral cortex development from human embryonic stem cells. Neuron 83, 51–68. 10.1016/j.neuron.2014.05.013.

44. Traxler, L., Lagerwall, J., Eichhorner, S., Stefanoni, D., D’Alessandro, A., and Mertens, J. (2021). Metabolism navigates neural cell fate in development, aging and neurodegeneration. DMM Disease Models and Mechanisms 14. 10.1242/DMM.048993/271173.

45. Baumholtz, A.I., Simard, A., Nikolopoulou, E., Oosenbrug, M., Collins, M.M., Piontek, A., Krause, G., Piontek, J., Greene, N.D.E., and Ryan, A.K. (2017). Claudins are essential for cell shape changes and convergent extension movements during neural tube closure. Dev Biol 428, 25–38. 10.1016/J.YDBIO.2017.05.013.

46. Punovuori, K., Malaguti, M., and Lowell, S. (2021). Cadherins in early neural development. Cellular and Molecular Life Sciences 78, 4435–4450. 10.1007/S00018-021-03815-9/FIGURES/5.

47. Sasai, N., Kadoya, M., and Ong Lee Chen, A. (2021). Neural induction: Historical views and application to pluripotent stem cells. Dev Growth Differ 63, 26–37. 10.1111/DGD.12703.

48. Rosebrock, D., Arora, S., Mutukula, N., Volkman, R., Gralinska, E., Balaskas, A., Aragonés Hernández, A., Buschow, R., Brändl, B., Müller, F.J., et al. (2022). Enhanced cortical neural stem cell identity through short SMAD and WNT inhibition in human cerebral organoids facilitates emergence of outer radial glial cells. Nature Cell Biology 2022 24:6 24, 981–995. 10.1038/s41556-022-00929-5.

49. Cederquist, G.Y., Asciolla, J.J., Tchieu, J., Walsh, R.M., Cornacchia, D., Resh, M.D., and Studer, L. (2019). Specification of positional identity in forebrain organoids. Nature Biotechnology 2019 37:4 37, 436–444. 10.1038/s41587-019-0085-3.

50. Kadoshima, T., Sakaguchi, H., Nakano, T., Soen, M., Ando, S., Eiraku, M., and Sasai, Y. (2013). Self-organization of axial polarity, inside-out layer pattern, and species-specific progenitor dynamics in human ES cell-derived neocortex. Proc Natl Acad Sci U S A 110, 20284–20289. 10.1073/PNAS.1315710110/SUPPL_FILE/SM03.MOV.

51. Metzis, V., Steinhauser, S., Pakanavicius, E., Gouti, M., Stamataki, D., Ivanovitch, K., Watson, T., Rayon, T., Mousavy Gharavy, S.N., Lovell-Badge, R., et al. (2018). Nervous System Regionalization Entails Axial Allocation before Neural Differentiation. Cell. 10.1016/j.cell.2018.09.040.

52. Gouti, M., Tsakiridis, A., Wymeersch, F.J., Huang, Y., Kleinjung, J., Wilson, V., and Briscoe, J. (2014). In Vitro Generation of Neuromesodermal Progenitors Reveals Distinct Roles for Wnt Signalling in the Specification of Spinal Cord and Paraxial Mesoderm Identity. PLoS Biol 12, e1001937. 10.1371/JOURNAL.PBIO.1001937.

53. Rayon, T., Stamataki, D., Perez-Carrasco, R., Garcia-Perez, L., Barrington, C., Melchionda, M., Exelby, K., Lazaro, J., Tybulewicz, V.L.J., Fisher, E.M.C., et al. (2020). Species-specific pace of development is associated with differences in protein stability. Science (1979) 369. 10.1126/SCIENCE.ABA7667/SUPPL_FILE/ABA7667_TABLES2.XLSX.

54. Greig, L.C., Woodworth, M.B., Galazo, M.J., Padmanabhan, H., and Macklis, J.D. (2013). Molecular logic of neocortical projection neuron specification, development and diversity. Nature Reviews Neuroscience 2013 14:11 14, 755–769. 10.1038/nrn3586.

55. Arlotta, P., Molyneaux, B.J., Jabaudon, D., Yoshida, Y., and Macklis, J.D. (2008). Ctip2 Controls the Differentiation of Medium Spiny Neurons and the Establishment of the Cellular Architecture of the Striatum. Journal of Neuroscience 28, 622–632. 10.1523/JNEUROSCI.2986-07.2008.

56. Ianevski, A., Giri, A.K., and Aittokallio, T. (2022). Fully-automated and ultra-fast cell-type identification using specific marker combinations from single-cell transcriptomic data. Nature Communications 2022 13:1 13, 1–10. 10.1038/s41467-022-28803-w.

57. Gagliano, O., Luni, C., Qin, W., Bertin, E., Torchio, E., Galvanin, S., Urciuolo, A., and Elvassore, N. (2019). Microfluidic reprogramming to pluripotency of human somatic cells. Nat Protoc, 722–737. 10.1038/s41596-018-0108-4.

58. de Sousa, J.A., Wong, C.W., Dunkel, I., Owens, T., Voigt, P., Hodgson, A., Baker, D., Schulz, E.G., Reik, W., Smith, A., et al. (2023). Epigenetic dynamics during capacitation of naïve human pluripotent stem cells. Sci Adv 9. 10.1126/SCIADV.ADG1936/SUPPL_FILE/SCIADV.ADG1936_TABLES_S1_TO_S18.ZIP.

59. Osnato, A., Brown, S., Krueger, C., Andrews, S., Collier, A.J., Nakanoh, S., Londoño, M.Q., Wesley, B.T., Muraro, D., Brumm, A.S., et al. (2021). Tgfβ signalling is required to maintain pluripotency of human naïve pluripotent stem cells. Elife 10. 10.7554/ELIFE.67259.

60. Guo, G., Stirparo, G.G., Strawbridge, S.E., Spindlow, D., Yang, J., Clarke, J., Dattani, A., Yanagida, A., Li, M.A., Myers, S., et al. (2021). Human naive epiblast cells possess unrestricted lineage potential. Cell Stem Cell 28, 1040–1056.e6. 10.1016/j.stem.2021.02.025.

61. Bayerl, J., Ayyash, M., Shani, T., Manor, Y.S., Gafni, O., Massarwa, R., Kalma, Y., Aguilera-Castrejon, A., Zerbib, M., Amir, H., et al. (2021). Principles of signaling pathway modulation for enhancing human naive pluripotency induction. Cell Stem Cell 28, 1549–1565.e12. 10.1016/J.STEM.2021.04.001/ATTACHMENT/87C21D9C-6CAD-4C27-977E-D23640A61FB0/MMC9.PDF.

62. Kim, E.J.Y., Sorokin, L., and Hiiragi, T. (2022). ECM-integrin signalling instructs cellular position sensing to pattern the early mouse embryo. Development (Cambridge) 149. 10.1242/DEV.200140/273721/AM/ECM-INTEGRIN-SIGNALLING-INSTRUCTS-CELLULAR.

63. Jain, A., Gut, G., Sanchis-Calleja, F., Okamoto, R., Streib, S., He, Z., Zenk, F., Santel, M., Seimiya, M., Holtackers, R., et al. (2023). Morphodynamics of human early brain organoid development. bioRxiv, 2023.08.21.553827. 10.1101/2023.08.21.553827.

64. Chiaradia, I., Imaz-Rosshandler, I., Nilges, B.S., Boulanger, J., Pellegrini, L., Das, R., Kashikar, N.D., and Lancaster, M.A. (2023). Tissue morphology influences the temporal program of human brain organoid development. Cell Stem Cell 30, 1351–1367.e10. 10.1016/j.stem.2023.09.003.

65. Yamaguchi, T.P. (2001). Heads or tails: Wnts and anterior–posterior patterning. Current Biology 11, R713–R724. 10.1016/S0960-9822(01)00417-1.

66. Velasco, S., Kedaigle, A.J., Simmons, S.K., Nash, A., Rocha, M., Quadrato, G., Paulsen, B., Nguyen, L., Adiconis, X., Regev, A., et al. (2019). Individual brain organoids reproducibly form cell diversity of the human cerebral cortex. Nature 2019 570:7762 570, 523–527. 10.1038/s41586-019-1289-x.

67. Tremblay, R., Lee, S., and Rudy, B. (2016). GABAergic Interneurons in the Neocortex: From Cellular Properties to Circuits. Neuron 91, 260–292. 10.1016/J.NEURON.2016.06.033/ASSET/24A0360E-A503-4B14-A432-9BB4BC681BD8/MAIN.ASSETS/GR7.JPG.

68. Krienen, F.M., Goldman, M., Zhang, Q., C. H. del Rosario, R., Florio, M., Machold, R., Saunders, A., Levandowski, K., Zaniewski, H., Schuman, B., et al. (2020). Innovations present in the primate interneuron repertoire. Nature 2020 586:7828 586, 262–269. 10.1038/s41586-020-2781-z.

69. Yu, Y., Zeng, Z., Xie, D., Chen, R., Sha, Y., Huang, S., Cai, W., Chen, W., Li, W., Ke, R., et al. (2021). Interneuron origin and molecular diversity in the human fetal brain. Nature Neuroscience 2021 24:12 24, 1745–1756. 10.1038/s41593-021-00940-3.

70. Sutherland, G.R., Gedeon, A., Kornman, L., Donnelly, A., Byard, R.W., Mulley, J.C., Kremer, E., Lynch, M., Pritchard, M., Yu, S., et al. (1991). Prenatal Diagnosis of Fragile X Syndrome by Direct Detection of the Unstable DNA Sequence. New England Journal of Medicine 325, 1720–1722. 10.1056/NEJM199112123252407/ASSET/176DB420-705D-448F-A02A-744D6C5F1333/ASSETS/IMAGES/LARGE/NEJM199112123252407_F3.JPG.

71. Dockendorff, T.C., and Labrador, M. (2019). The Fragile X Protein and Genome Function. Mol Neurobiol 56, 711–721. 10.1007/S12035-018-1122-9/FIGURES/4.

72. Bardoni, B., Mandel, J.L., and Fisch, G.S. (2000). FMR1 gene and fragile X syndrome. American Journal of Medical Genetics - Seminars in Medical Genetics 97, 153–163. 10.1002/1096-8628(200022)97:2<153::AID-AJMG7>3.0.CO;2-M.

73. Qin, D., Xia, Y., and Whitesides, G.M. (2010). Soft lithography for micro- and nanoscale patterning. Nature Protocols 2010 5:3 5, 491–502. 10.1038/nprot.2009.234.

74. Bredenkamp, N., Yang, J., Clarke, J., Stirparo, G.G., von Meyenn, F., Dietmann, S., Baker, D., Drummond, R., Ren, Y., Li, D., et al. (2019). Wnt Inhibition Facilitates RNA-Mediated Reprogramming of Human Somatic Cells to Naive Pluripotency. Stem Cell Reports. 10.1016/J.STEMCR.2019.10.009.

75. Krammer, T., Stuart, H.T., Gromberg, E., Ishihara, K., Cislo, D., Melchionda, M., Becerril Perez, F., Wang, J., Costantini, E., Lehr, S., et al. (2024). Mouse neural tube organoids self-organize floorplate through BMP-mediated cluster competition. Dev Cell 59, 1940–1953.e10. 10.1016/J.DEVCEL.2024.04.021/ATTACHMENT/DA1AE823-3ED7-408C-87B2-ADCDEFC803ED/MMC7.PDF.

76. Kinoshita, M., Barber, M., Mansfield, W., Cui, Y., Spindlow, D., Stirparo, G.G., Dietmann, S., Nichols, J., and Smith, A. (2021). Capture of Mouse and Human Stem Cells with Features of Formative Pluripotency. Cell Stem Cell 28, 453–471.e8. 10.1016/J.STEM.2020.11.005/ATTACHMENT/C2F8D6F0-5571-467C-9A34-B3C48520C983/MMC4.XLSX.

77. Zeng, B., Liu, Z., Lu, Y., Zhong, S., Qin, S., Huang, L., Zeng, Y., Li, Z., Dong, H., Shi, Y., et al. (2023). The single-cell and spatial transcriptional landscape of human gastrulation and early brain development. Cell Stem Cell 30, 851–866.e7. 10.1016/j.stem.2023.04.016.

78. Rifes, P., Isaksson, M., Rathore, G.S., Aldrin-Kirk, P., Møller, O.K., Barzaghi, G., Lee, J., Egerod, K.L., Rausch, D.M., Parmar, M., et al. (2020). Modeling neural tube development by differentiation of human embryonic stem cells in a microfluidic WNT gradient. Nature Biotechnology 2020 38:11 38, 1265–1273. 10.1038/s41587-020-0525-0.

